# ACSS2 mediates prenatal alcohol exposure-related morphological and behavioral phenotypes

**DOI:** 10.1101/2025.11.26.690774

**Authors:** KM Dodson, EM Periandri, A Yadav, M Lopes, AJ Barfield, A Ola, FN de Luna Vitorino, C Cearlock, BA Garcia, C Hill, SE Maloney, G Egervari

## Abstract

The metabolic enzyme Acetyl-CoA Synthetase 2 (ACSS2) recently emerged as an unexpected regulator of molecular and behavioral changes associated with alcohol use. Its role during prenatal exposure, however, remains unknown. Here, we use a combination of proteomic, genomic and behavioral approaches to establish ACSS2 as a key mediator of prenatal alcohol exposure-related phenotypes. We define the developmental window during which ACSS2 translocates to nuclei in the mouse brain, and show that alcohol-derived acetate is incorporated into fetal brain histone acetylation in utero. Using genetically engineered mice not expressing ACSS2, we demonstrate that loss of this enzyme attenuates chronic prenatal alcohol exposure-induced craniofacial abnormalities, motor function deficits, cognitive impairments as well as associated chromatin and gene expression changes in the dorsal hippocampus and the cerebellar vermis. Our results outline a previously unknown mechanism underlying prenatal alcohol exposure-related phenotypes regulated by ACSS2, which will inform the development of future therapeutic interventions.

**HIGHLIGHTS:**

- ACSS2 translocates to nuclei during in utero brain development
- Alcohol-derived acetate is incorporated into fetal brain histone acetylation
- Prenatal alcohol exposure results in long-lasting and ACSS2-dependent chromatin and gene expression changes in the hippocampus and cerebellar vermis
- Loss of ACSS2 attenuates molecular changes, craniofacial abnormalities and cognitive impairments linked to prenatal alcohol exposure

## INTRODUCTION

Despite its known teratogenic properties, ethanol consumption during pregnancy is widespread. A recent study found that approximately 1 in 7 pregnant adults consumed alcoholic beverages within the past 30 days, and 1 in 20 reported binge drinking.^1^ As a result, Fetal Alcohol Spectrum Disorder (FASD) remains prevalent: An estimated 2-5% of school age children may have FASD in the United States.^2^ Moreover, as FASD is often mischaracterized as other neurodevelopmental disorders such as attention-deficit/hyperactivity disorder (ADHD), autism spectrum disorder (ASD) or other learning disabilities, the real number of affected children is potentially even higher.^3^

While symptoms and severity vary among individuals, FASD is generally characterized by craniofacial abnormalities,^4^ irregular brain development,^5^ and cognitive or neurodevelopmental impairments that range from memory deficits to seizures.^6,7^ Although the spectrum of symptoms that result from prenatal alcohol exposure (PAE) is well documented, the complex etiology of these disorders remains poorly understood, underscoring the need to explore novel mechanisms that might contribute to FASD. Recent studies suggested that cognitive impairments associated with PAE may be the result of epigenetic modifications including DNA methylation and histone acetylation.^8,9^ While histone acetylation, which enables an active state of transcription, has been tightly linked to both memory formation and postnatal alcohol exposure,^10–12^ impairments of this epigenetic mechanism have not been widely investigated in PAE paradigms.

A growing body of evidence indicates that epigenetic modifications may be driven by metabolites and metabolic enzyme.^13–16^ For instance, we recently found that following acute ethanol exposure, exogenous alcohol-derived acetate is incorporated into histone acetylation.^17^ This incorporation is mediated by Acetyl-CoA Synthetase 2 (ACSS2), a metabolic enzyme that converts acetate into acetyl-CoA. Acetyl-CoA is the primary substrate of histone acetyl transferase proteins (HATs) and is necessary for histone acetylation.^18^ During alcohol exposure in adult mice, ACSS2 mediates epigenetic and transcriptional outcomes and is required for the formation of alcohol-associated memories. Indeed, loss of ACSS2 results in significantly decreased voluntary alcohol intake.^17,19^ Despite emerging as a key neuroactive metabolic enzyme in the context of adult alcohol exposure, the role of ACSS2 during PAE remains unknown.

Here, we define the developmental window during which ACSS2 becomes nuclear localized in the brain and show that alcohol-derived acetate is incorporated into fetal brain histone acetylation at different developmental timepoints. Further, we combine advanced proteomic, transcriptional, imaging and behavioral approaches in the context of prenatal alcohol exposure to establish ACSS2 as a critical regulator of FASD-related phenotypes. We show that loss of ACSS2 attenuates craniofacial abnormalities and cognitive impairments and characterize the underlying chromatin and transcriptional changes. Overall, our study outlines a previously unknown aspect of FASD etiology driven by ACSS2-mediated metabolic-epigenetic interactions, which may inform the development of novel therapeutic interventions in the future.

## RESULTS

### ACSS2 translocates to nuclei during in utero mouse brain development

To assess when ACSS2 translocates into the nucleus during fetal development, we sacrificed saline- or ethanol-injected pregnant dams and extracted fetal brains at embryonic days E10.5, E11.5, and E17.5 (**Figure 1A**). These developmental timepoints represent fundamental events during neurodevelopment such as the completion of the neural tube at E10.5,^20^ peak neurogenesis at E11.5,^21,22^ and the final phase of in utero development at E17.5^23^. To determine the subcellular localization of ACSS2, we performed subcellular fractionation in conjunction with Western blotting in fetal brains at the selected timepoints (**Figure 1B**).

**Figure 1:**
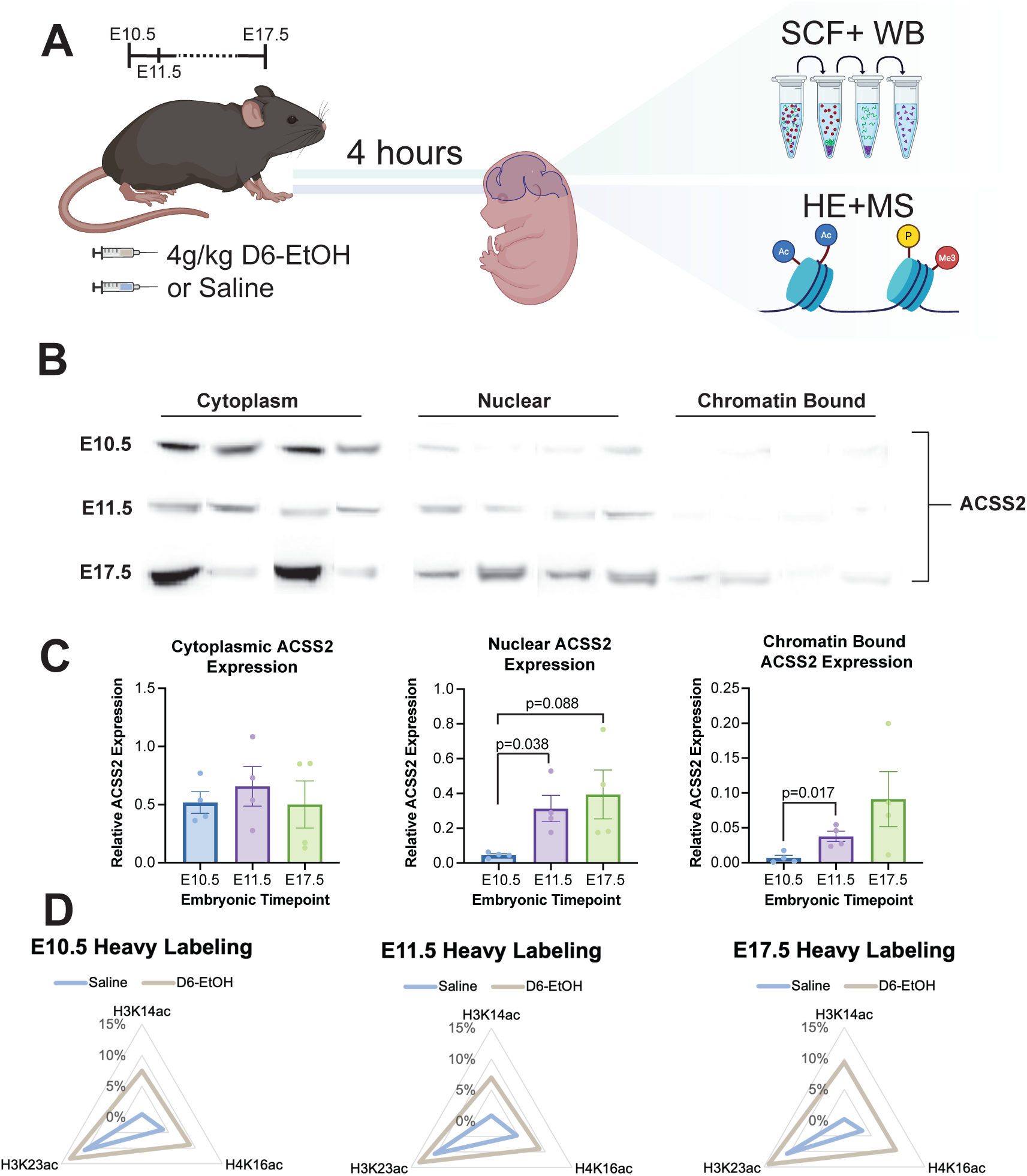
ACSS2 localization and alcohol-induced histone acetylation during in utero mouse brain development. **(A)** Schematic outlining fetal alcohol exposure, and downstream molecular assays including subcellular fraction with western blot and histone extraction with liquid chromatography-tandem mass spectrometry. **(B)** Western blot of ACSS2 expression in cytoplasmic, nuclear, and chromatin bound fractions. Sample size is n=4 fetal brain pools per timepoint. **(C)** Relative ACSS2 expression at embryonic days (E)10.5, E11.5, and E17.5 in cytoplasmic (left), nuclear (middle), and chromatin-bound (right) fractions. Protein expression is normalized to total histone proteins per sample. Sample Size is n=4 fetal brain pools per timepoint. **(D)** Radar charts showing incorporation of heavy-labeled acetyl marks into the fetal brain after D6-EtOH or saline exposure in the E10.5 (left), E11.5 (middle), and E17.5 (right) fetal brain.

We found consistent expression of ACSS2 in cytoplasmic fractions at all developmental timepoints irrespective of treatment. However, nuclear levels of ACSS2 tended to be lower at E10.5 compared to E11.5 and E17.5 (Welch’s t-test: [E10.5 vs E11.5] t_3.062_=3.509, *p*=0.0380; Welch’s t-test: [E10.5 vs E17.5] t_3_=2.485, *p*=0.0884) (**Figure 1C**), suggesting that ACSS2 translocates to nuclei around this developmental stage. Similarly, we observed higher levels of ACSS2 in the chromatin-bound fraction at E11.5 compared to E10.5 (Welch’s t-test: [E10.5 vs E11.5] t_4_=3.706 *p*=0.017).

Together, our results indicate that ACSS2 translocates to nuclei during fetal brain development, near the time of peak neurogenesis at E11.5. This timing aligns with the emerging role of ACSS2 in neuronal differentiation and function.^24^

### Alcohol-derived acetate is incorporated into histone acetylation in the developing brain

We have previously shown that alcohol-derived acetate is incorporated into histone acetylation in the near-term fetal brain when pregnant dams are exposed to alcohol at E18.5.^17^ Given our current findings that ACSS2 is expressed earlier in development, we hypothesized that developing brains might be susceptible to alcohol-driven histone acetylation throughout pregnancy. To test this, pregnant dams were exposed to either 4 g/kg deuterated D6-ethanol (D6-EtOH) or equivalent volume of saline i.p. at the same three developmental timepoints (**Figure 1A**). We performed histone extraction followed by stable isotope labeling liquid chromatography tandem mass spectrometry (LC-MS/MS) of fetal brains and quantified the of heavy labeling of three abundant and thus reliably detected acetylated histone residues: H3K14ac, H3K23ac and H4K16ac.

We found increased heavy labeling of all three acetylated histone peptides following D6-EtOH administration at all three developmental timepoints (H3K23ac two-way ANOVA: [treatment] F_1,18_=234, *p*<0.0001; H4K16ac two-way ANOVA: [treatment] F_1,18_=331, *p*<0.0001; H3K14ac two-way ANOVA: [treatment], F_1,18_=478, *p*<0.0001, interaction F_2,18_=7, *p*=0.0054) (**Figure 1D**), suggesting that fetal ACSS2 expression is sufficient to enable acetate incorporation. Interestingly, H3K14ac showed a timepoint-specific effect, with E17.5 displaying significantly higher levels of heavy-label incorporation following D6-EtOH exposure compared to E10.5 and E11.5 (H3k14ac two-way ANOVA interaction: [timepoint × treatment] F_2,18_=7.0, *p*=0.0054; post hoc Šidák: D6-EtOH [E10.5 vs E17.5] t_18_=3.225, adj. *p*=0.0279; D6-EtOH [E11.5 vs E17.5] t_18_=4.030, adj. *p*=0.0047) (**Figure S1A)**.

Overall, our stable isotope labeling LC-MS/MS results revealed that, similarly to adult and near-term mice, alcohol-derived acetate is incorporated into histone acetylation in the developing brain in utero. While cytoplasmic ACSS2 expression appears to be sufficient to drive this pathway, the translocation to ACSS2 to the nucleus and chromatin might increase the extent of acetyl-group deposition on specific histone lysine residues.

### In utero alcohol exposure leads to long-lasting hippocampal and cerebellar chromatin remodeling in an ACSS2-dependent manner

Despite FASD being frequently diagnosed in adolescence, existing research primarily focused on short-term epigenetic and transcriptional outcomes, while the long-lasting gene regulatory consequences of prenatal ethanol exposure remain poorly understood. To address this critical knowledge gap, we exposed wildtype (WT) and ACSS2 knockout (ACSS2^KO^) mice to chronic prenatal alcohol exposure (cPAE) or chronic prenatal saline exposure (cPSE) and investigated chromatin and gene expression in the adolescent offspring at approximately postnatal day (PND) 45 (**Figure 2A**). In light of our observations that alcohol-derived acetate incorporation occurs in fetal brains at both early and late developmental timepoints, we utilized a model of exposure lasting the entire duration of pregnancy. We focused on two key brain regions: 1. the cerebellar vermis based on prior reports of vermis-mediated motor deficits associated with prenatal alcohol exposure in both humans and rodents;^25,26^ and 2. the dorsal hippocampus (dHPC) which is impacted by alcohol-derived acetate^17^ and regulates long-term memory, impairments of which are hallmarks of FASD.^27^

**Figure 2:**
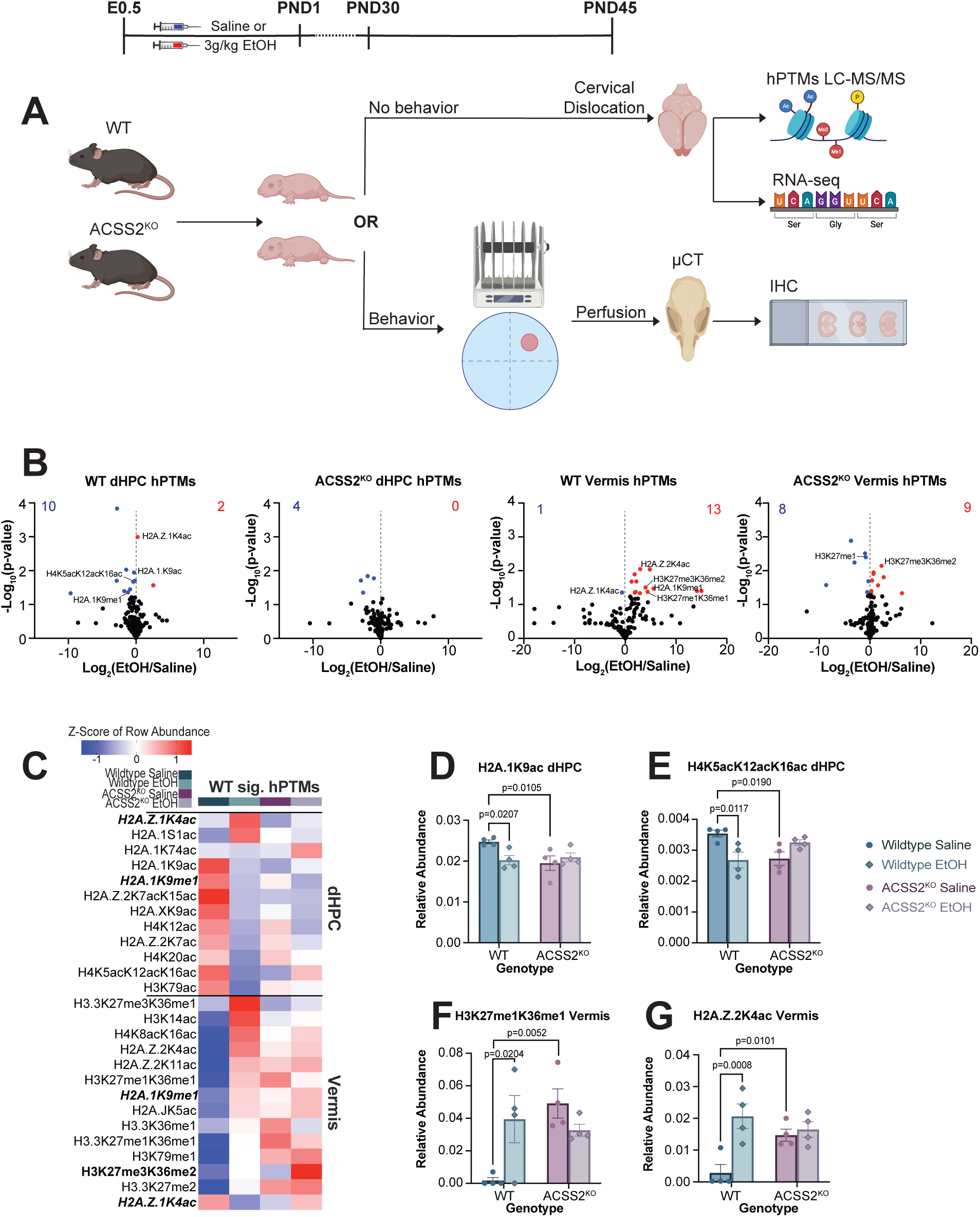
Chronic prenatal alcohol exposure affects histone post translational modifications in a brain region and genotype specific manner. **(A)** Schematic of the chronic prenatal alcohol exposure (cPAE) or prenatal saline exposure (cPSE) paradigm, including behavioral assays (Morris water maze/accelerating rotarod), imaging (µCT/immunohistochemistry) and molecular techniques (RNA sequencing/ histone extraction with liquid chromatography-tandem mass spectrometry). **(B)** Volcano plots depicting differentially expressed hPTMs for WT vermis (left), WT dorsal hippocampus (middle left), ACSS2^KO^ vermis (middle right), and ACSS2^KO^ dorsal hippocampus (right). **(C)** Heatmap of average row Z-score of relative abundance for significant WT hPTMs sorted from most upregulated to most downregulated following cPAE in the dHPC (upper) and vermis (lower). Overlapping hPTMs are bolded. **(D-G.)** Bar charts depicting the average relative abundance of hPTMs in dHPC or vermis following cPAE or cPSE in WT and ACSS2^KO^ mice. **(D)** Average relative abundance of H2A.1K9ac in the dHPC. **(E)** Average relative abundance of H4K5acK12acK16ac in the dHPC. **(F)** Average relative abundance of H3K27me1K36me1 in the vermis. **(G)** Average relative abundance of H2A.Z.2K4ac in the vermis. Samples sizes are n=4 per genotype per condition.

We used LC-MS/MS to examine histone post-translational modifications (hPTMs) in WT and ACSS2^KO^ dHPC and vermis comparing mice that underwent cPAE or cPSE. In the WT dHPC, we found preferential downregulation of hPTMs (10 downregulated peptides, 2 upregulated peptides). The ACSS2^KO^ dHPC displayed a similar, but muted pattern (4 downregulated peptides, 0 upregulated peptides). In contrast, the WT vermis exhibited largely upregulated hPTMs (1 downregulated peptide, 13 upregulated peptides) while the ACSS2^KO^ vermis showed nearly equal up- and downregulation of hPTMs following cPAE (8 downregulated peptides, 9 upregulated peptides) (**Figure 2B**). Across genotypes or regions, there were only three overlapping hPTMs affected by cPAE. Two of these, H2A.1K9me1and H2A.Z.1K4ac, were significantly affected in the WT dHPC and vermis, both displaying opposite directionality following cPAE. H2A.Z.1K4ac, which we previously found to be significantly affected in the dHPC following acetate injection,^28^ was the most significantly upregulated hPTM in the WT dHPC (**Figure 2C, top**) and the most significantly downregulated hPTM in the WT vermis (**Figure 2C, bottom**). Lastly, H3K27me3K36me2 was downregulated following cPAE in the vermis regardless of genotype (**Figure 2C & S1A**).

Strikingly, 67% of all significantly affected hPTMs in WT mice (dHPC and vermis) were acetylated H2A or H2A.Z peptides. In ACSS2^KO^ mice, we saw a reduced effect of cPAE on these peptides, with 43% of affected hPTMs relating to H2A or H2A.Z (**Figure S1A**). Interestingly, acetylation of the histone variant H2A.Z.2 was dysregulated more often in WT mice following cPAE, while H2A.Z.1 acetylation was more commonly affected in the ACSS2^KO^ brain. In general, acetyl peptides comprised 90% of all significantly affected hPTMs in the WT dHPC, the majority of which were decreased following cPAE (**Figure 2C**). Many of these epigenetic changes were specific to genotype. For example, H2A.1K9ac was significantly decreased in the dHPC of WT, but not ACSS2^KO^ mice following cPAE (Two-way ANOVA interaction: [genotype × treatment] F_1,12_=6.072 *p*=0.0298; post hoc Šidák: WT [cPSE vs cPAE] t_12_=2.661, adj. *p*=0.0207; ACSS2^KO^ [cPSE vs cPAE] t_12_=0.7566, adj. *p*=0.7126) (**Figure 2D**). H4K5acK12acK16ac was decreased in the dHPC of WT mice, but not in the dHPC of ACSS2^KO^ mice following cPAE (two-way ANOVA interaction: [genotype × treatment] F_1,12_=14, *p*=0.0025; post hoc Šidák: WT [cPSE vs cPAE] adj. *p*=0.0058; ACSS2^KO^[cPSE vs cPAE] t_12_=3.344, adj. *p*=0.1255) (**Figure 2E**).

In the vermis, cPAE induced genotype-specific mono-methylation of several histone peptides (**Figure 2C**). For example, H3K27me1K36me1 was upregulated in the WT vermis but displayed no significant effect in the ACSS2^KO^ vermis following cPAE (two-way ANOVA interaction: [genotype × treatment] F_1,12_ =9.5, *p*=0.0094; post hoc Sidak WT [cPSE vs cPAE] t_12_ =3.041, adj. *p*=0.0204; ACSS2^KO^[cPSE vs cPAE] t_12_=0.3736, adj. *p*=1.329) (**Figure 2F**). Further, as an example of WT-specific H2A.Z.2ac dysregulation, H2A.Z.2K4ac also exhibited a genotype-specific response to cPAE, where it increased in the WT vermis, but displayed no change in ACSS2^KO^ mice (two-way ANOVA interaction: [genotype × treatment] F_1,12_=8.0, *p*=0.0148; post hoc Sidak: WT [cPSE vs cPAE] t_12_ =4.47, adj. *p*=0.0015; ACSS2^KO^[cSPE vs cPAE] t_12_=0.4485, adj. *p*=0.6618)(**Figure 2G)**.

Overall, our findings indicate that cPAE results in long-lasting hPTM changes in key brain regions associated with FASD, with a particularly strong impact on histone H2A and its variants. Importantly, the majority of these effects are genotype-specific, emphasizing the key role of ACSS2 in alcohol-induced chromatin remodeling in the context of PAE.

### cPAE results in long-lasting gene expression changes in the dHPC and vermis in an ACSS2-dependent manner

We used RNA sequencing to explore the lasting transcriptomic impact of cPAE in the dHPC and vermis of adolescent WT and ACSS2^KO^ mice. Gene level analyses revealed 248 differentially expressed genes (DEGs) (104 upregulated and 144 downregulated) in the dHPC of WT cPAE mice compared to cPSE controls (**Figure 3A, top left**). Despite sufficient set sizes, gene ontology (GO) analysis identified no significantly enriched terms among dHPC DEGs, suggesting that while cPAE had a marked effect on gene expression, transcriptional changes were not consistently correlated with specific functions. We next used STRING to construct protein-protein interaction (PPI) networks related to DEGs and observed 2 clusters (**Figure S2A**. Cluster 1 included genes related to circadian regulation (e.g., *Per2/3* and *Cry1* upregulated) and synaptic activity (e.g., *Egr2, Fos, Npas4, Npy2r,* and *Ptgs2* downregulated), whereas cluster 2 was characterized by downregulation of genes involved in ribosome biogenesis (RPLs) and mitochondrial function (e.g., *Chchd2, Cox17, Ndufa4/5, Timm10*).

**Figure 3.**
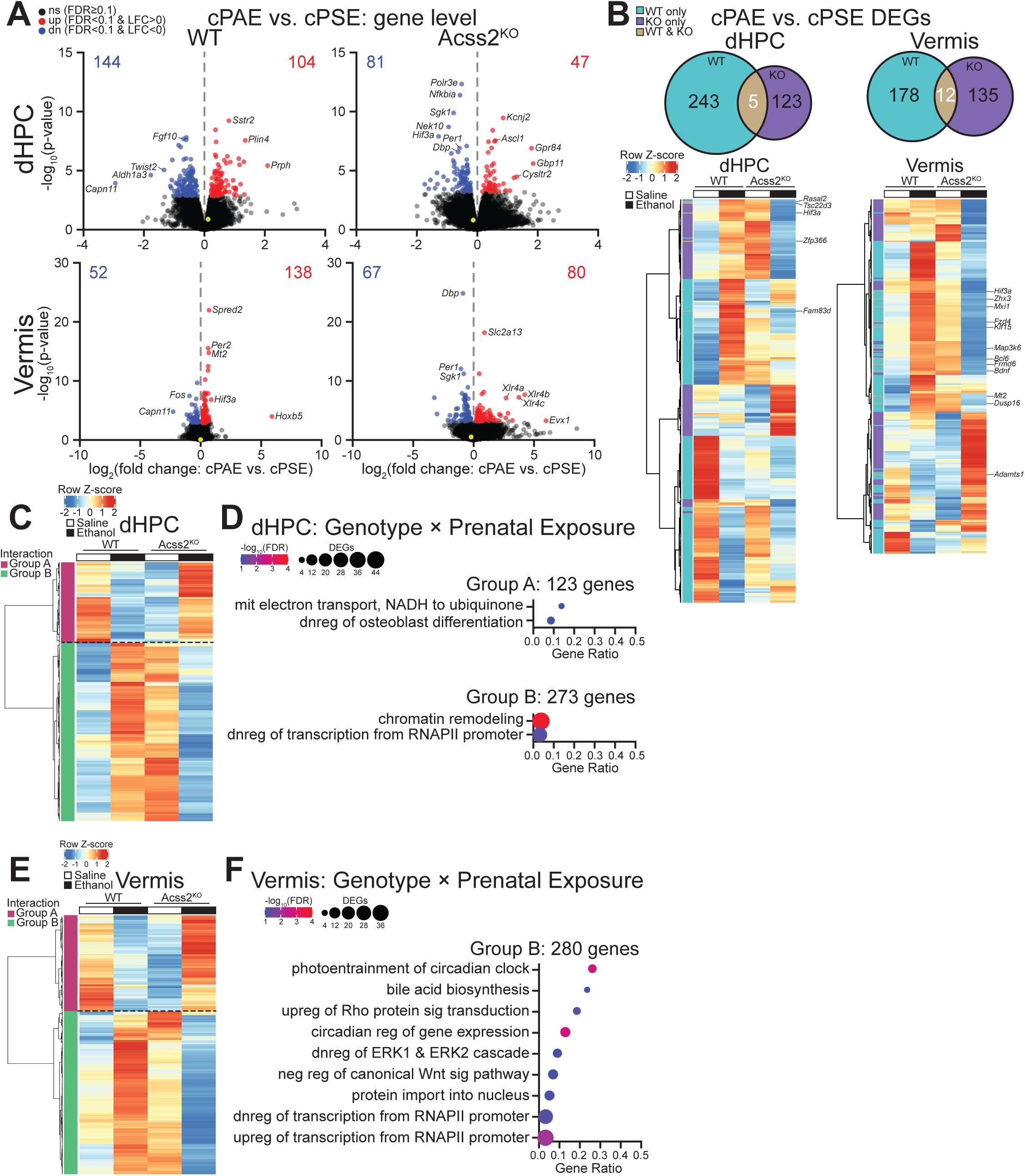
Chronic prenatal alcohol exposure drives lasting gene expression changes in the dorsal hippocampus and vermis in an ACSS2-dependent manner. **(A)** Volcano plots showing differential gene expression in the dHPC (upper) and vermis (lower) of young adult WT (left) and ACSS2^KO^ (right) mice subjected to cPAE vs. respective cPSE controls. Dots represent genes (black not significant, red significantly upregulated, blue significantly downregulated, yellow *Acss2*), and numbers show quantity of DEGs. **(B)** Venn diagrams (upper) showing overlap of DEGs between genotypes within each region and heatmaps (lower) showing average row Z-score of VST-normalized gene counts of DEGs. **(C)** Heatmap of genes with significant ACSS2 genotype × prenatal exposure interaction in the dHPC. **(D)** Significantly enriched GO terms for significant interaction genes in the dHPC. **(E)** Heatmap of significant interaction genes in the vermis. **(F)** Significantly enriched GO terms for significant interaction genes in the vermis. For dot plots of enriched GO terms, dot color represents -log_10_(FDR), dot size represents gene count (number of DEGs within a term), and gene ratio is the quantity of DEGs out of detected annotated genes for a term. n=4 mice per genotype per condition. Abbreviations: dnreg, downregulation; mit, mitochondrial; neg reg, negative regulation; reg, regulation; upreg, upregulation.

In the dHPC of ACSS2^KO^ mice, cPAE resulted in 128 DEGs (47 upregulated and 81 downregulated) (**Figure 3A, top right**), which were significantly enriched for GO terms related to downregulation of neuron differentiation and upregulation of transcription from RNA polymerase II (RNAPII) promoter (**Figure S2B**). The PPI network of DEGs in the ACSS2^KO^ dHPC contained one large cluster (**Figure S2C**), which included genes related to downregulation of circadian regulation (e.g., *Per1, Ciart, Dbp, Nr1d1*) and neuronal stress response genes (e.g., *Foxo3, Ddit4, Nfkbia, Sgk1*), and upregulation of genes associated with neurogenesis (e.g., *Sox2/9, Olig2, Id2, Ascl)* and protein folding (e.g., *HSpa5/90b1, Calr, Pdia6*). Of note, there was limited overlap between WT and ACSS2^KO^ DEGs in the dHPC (5), all of which showed opposite directionality (**Figure 3B**).

In the vermis of WT mice, cPAE resulted in 190 DEGs (138 upregulated and 52 downregulated) (**Figure 3A, bottom left**), with GO terms significantly enriched for positive regulation of Hippo signaling and response to corticosterone (**Figure S2D, upper**). The PPI network identified one cluster (**Figure S2E**) including upregulated genes related to transcriptional/epigenetic regulation (e.g., *Bcl6, Ets2, Foxo1/3, Klf9/15,* etc.) and synaptic function (e.g., *Bdnf, Crhr1, Nedd4l, Rtn4r/4rl1,* etc.), and downregulated genes related to reduced activity-dependent transcription (e.g., *Fos, Npas4, Dusp1*).

Finally, in the vermis of ACSS2^KO^ mice, we observed 147 DEGs (80 upregulated and 67 downregulated) following cPAE (**Figure 3A, bottom right**), which were significantly enriched for various biological processes such as dopamine secretion, upregulation of transcription from RNAPII promoter, and immune-related functions (e.g., negative regulation of dendritic cell apoptosis, regulation of B cell differentiation) (**Figure S2D, lower**). PPI network analysis revealed 2 clusters (**Figure S2F**). The larger cluster showed upregulation of genes associated with neurodevelopment (e.g., *Bmpr1b, Tgfbr2, Notch1,* etc.), and downregulation of genes associated with transcriptional regulation (e.g., *Bcl6, Klf2/15, Cebpd*), and neuronal stress response (*e.g., Bdnf, Nfkbia, Nr4a2/3, Ddit4).* The smaller cluster displayed downregulation of genes associated with circadian rhythm regulation (e.g., *Ciart, Per1/3, Nr1d1/2*). Similarly to the dHPC, we observed few (12) overlapping DEGs between the WT and KO vermis, all showing opposite directionality in the two genotypes (**Figure 3B**). This suggests that ACSS2 might play an important role in cPAE-induced transcriptional changes in both brain regions.

To further assess genotype-specific effects of cPAE, we next performed genotype × treatment interaction analysis. Within both regions we found hundreds of genes exhibiting significant interaction, which generally fell into 2 categories: group A genes were downregulated in WT but upregulated in ACSS2^KO^ cPAE mice, and group B genes were upregulated in WT but downregulated in ACSS2^KO^ cPAE mice. In the dHPC, we identified 396 genes with a significant interaction (123 in Group A, and 273 in Group B) (**Figure 3C and S3A**). GO analysis revealed that group A genes were significantly enriched for downregulation of osteoblast differentiation and mitochondrial electron transport, NADH to ubiquinone (**Figure 3D, upper**), whereas group B genes were enriched for chromatin remodeling and downregulation of transcription from RNAPII promoter (**Fig. 3D, lower**). In the vermis, 445 genes showed a significant genotype × treatment interaction (165 DEGs in Group A, and 280 DEGs in Group B) (**Figures 3E and S3A**). No GO terms were identified within Group A genes, however, Group B genes were enriched for various biological processes, such as protein import into nucleus, regulation of transcription from RNAPII promoter, and circadian-related functions (**Figure 3F**).

Taken together, we found that cPAE induces long-lasting transcriptional changes in the dHPC and vermis. Emphasizing the important role of ACSS2, loss of this enzyme markedly affected cPAE-associated transcriptional outcomes in both brain regions.

### Transcript-level analysis reveals long-lasting ACSS2-dependent isoform expression changes in the dHPC and vermis following cPAE

Gene level analysis can mask important isoform-specific transcriptional changes. In addition, alcohol exposure has been previously linked to impairments of alternative splicing.^29,30^ We thus performed transcript level analysis to test whether ACSS2 expression affects these outcomes in the context of cPAE.

We found dysregulated transcript expression in both brain regions following cPAE. In the dHPC, there were 757 differentially expressed transcripts (DETs) (229 upregulated from 83 genes and 528 downregulated from 178 genes) in WT mice and 524 DETs (131 upregulated from 57 genes and 393 downregulated from 146 genes) in ACSS2^KO^ mice (**Figure 4A, upper**). In the vermis, there were 1,044 DETs (647 upregulated from 233 genes and 397 downregulated from 67 genes) in WT mice and 394 DETs (167 upregulated from 75 genes and 227 downregulated from 87 genes) in ACSS2^KO^ mice (**Figure 4A, lower**). Regardless of brain region or genotype, the majority of DETs were previously identified as DEGs (**Figure 4B**), matching in directionality between gene and transcript levels.

**Figure 4.**
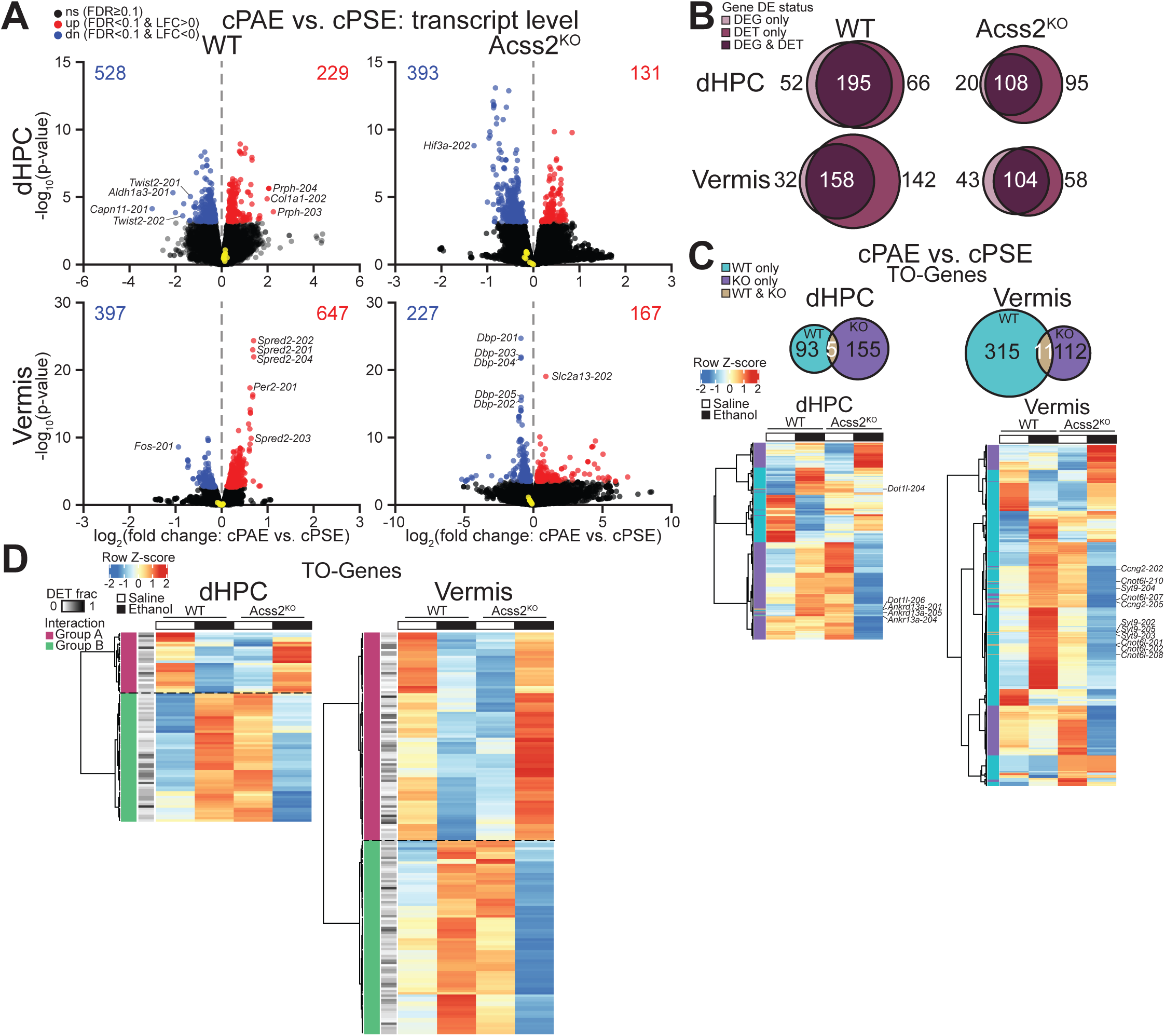
Chronic prenatal alcohol exposure drives lasting transcript expression changes in the dorsal hippocampus and vermis. **(A)** Volcano plots showing differential transcript expression in the dHPC (upper) and vermis (lower) of young adult WT (left) and ACSS2^KO^ (right) mice subjected to cPAE vs. respective cPSE controls. Dots represent transcripts (black not significant, red significantly upregulated, blue significantly downregulated, yellow encoded by *Acss2*), and numbers are quantity of DETs. **(B)** Venn diagrams showing overlap of DEGs and genes encoding DETs. **(C)** Venn diagrams (upper) showing overlap of TO-genes between genotypes within each region and heatmaps (lower) showing average row Z-score of VST-normalized transcript counts of TO-genes. **(D)** Heatmaps significant genotype × exposure interaction TO-genes in dHPC (left) and vermis (right), annotated with DET fraction (fraction of DETs of all transcript isoforms corresponding to a gene). n=4 mice per genotype per condition.

However, nearly a third of DETs were encoded by genes not identified as a DEG (**Figure 4B**), indicating isoform-specific effects of cPAE. PPI network analysis of DETs not associated with DEGs (transcript level only [TO-genes]), revealed 2 clusters in the WT dHPC, which included genes related to epigenetic regulation (e.g., *Dot1l, Dnmt3a,* and *Cbx3*) (**Figure S4A)**. In the ACSS2^KO^ dHPC, we observed 2 clusters including genes related to cellular signaling pathways *(e.g., Sdc4, Ptk2, Lamc1)* and downregulation of genes associated with transcriptional regulation (*e.g., Safb, Safb2, Spen, Snrnp70*) **(Figure S4B).** PPI network analysis of TO-genes identified two clusters in the WT vermis and none in ACSS2^KO^ vermis. In the WT vermis TO-gene clusters had biological functions associated with upregulated neuronal stress responses (e*.g, Manf, Eif2ak3, Map1lc3b, Ulk1, Ctsa*) and cellular signaling (*e.g., Arhgef7, Itgb5, Crk*), and downregulated genes involved in ribosomal synthesis (e*.g., Rpl13a/12, Rps23, Utp20*) **(Figure S4C).**

Interestingly, TO-genes shared between genotypes were limited both in the dHPC (5 overlapping) and in the vermis (11 overlapping) (**Figure 4C, upper**). Several of these shared transcripts were associated with genes previously implicated in substance use disorders including *Dot1l* (2 DETs) or *Ankrd13a* (3 DETs) in the dHPC, *Ccng2* (2 DETs), *Cnot6l* (5 DETs), or *Syt9* (4 DETs) in the vermis (**Figure 4C, lower**)^31–35^ all shared TO-genes showed opposite directionality between genotypes (**Figure 4C, lower**), emphasizing the distinct effects of cPAE in WT and KO mice.

We next performed genotype × treatment interaction analysis for transcripts in both brain regions. In line with what we observed on the gene level, we found that significant interactions fell into two categories: Group A DETs were upregulated in ACSS2^KO^ and downregulated in WT, while group B DETs were upregulated in WT and downregulated in ACSS2^KO^. In the dHPC, 1,088 transcripts from 374 genes exhibited a significant interaction (**Figure S4D, left**), the majority of which were associated with genes previously identified in the gene level interaction analysis (**Figure S4E, left**). 81 transcripts, however, were associated with genes that showed no interaction (**Figure 4D, left and S4E left**). In the vermis, we identified 1,366 transcripts (from 532 genes) showing a significant interaction (**Figure S4D, right**), over half of which were also identified on the gene level (**Figure S4E, right**). Similarly to the dHPC, a large number (172) of transcripts were not associated with interaction genes (**Figure 4D, right and S4E right**), emphasizing isoform-specific effects potentially masked in gene level analyses.

Overall, these results suggest that ACSS2 expression plays a critical role in lasting transcriptional outcomes following cPAE.

### cPAE reduces hippocampal thickness in the male brain irrespective of ACSS2 genotype

Structural changes in key brain regions, such as the neocortex and hippocampus, are important phenotypic hallmarks of FASD.^36,37^ To investigate whether ethanol-induced abnormalities in brain development depend on ACSS2, we exposed WT and ACSS2^KO^ mice to cPAE or cPSE and performed immunohistochemistry (IHC) in the adolescent offspring (**Figure 2A**). Given the known differences in hippocampal volume between males and females, all IHC data was stratified by sex.^38,39^

We assessed thickness of three hippocampal regions (**Figure S5A**): CA1, CA2 and the dentate gyrus (DG) lower blade. In males, cPAE treatment reduced thickness of CA1 (two-way ANOVA: [treatment] F_1,12_=6.6, p=0.0243) (**Figure 5A**), CA2 (two-way ANOVA: [treatment] F_1,12_=8.3, p=0.0136) (**Figure 5B**), and the DG lower blade (two-way ANOVA: [treatment] F_1,12_=5.4, p=0.00371) (**Figure 5C**) regions irrespective of genotype. In females, hippocampal thickness was not affected by cPAE in either WT or ACSS2^KO^ mice (**Figure S5B-5D**).

**Figure 5:**
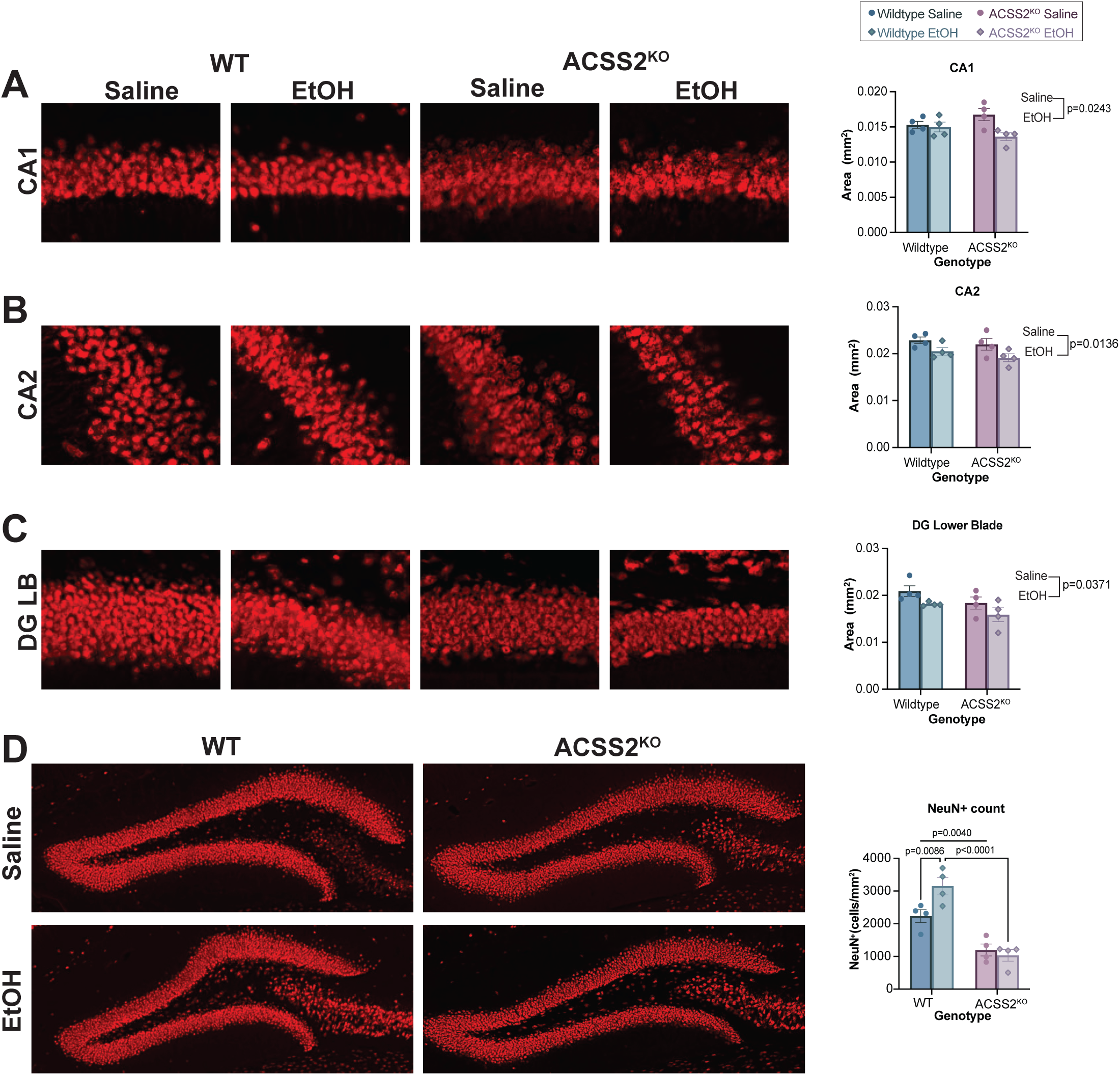
Chronic prenatal alcohol exposure reduces hippocampal thickness and dentate gyrus neuron count in the male brain. **(A)** NeuN+ staining of hippocampal region CA1 from left to right in WT cPSE, WT cPAE, ACSS2^KO^ cPSE, and ACSS2^KO^ cPAE and the average thickness of CA1 region (right most) in males. **(B)** NeuN+ staining of hippocampal region CA2 from left to right in WT cPSE, WT cPAE, ACSS2^KO^ cPSE, and ACSS2^KO^ cPAE and the average thickness of CA2 region (right most) in males. **(C)** NeuN+ staining of dentate gyrus lower blade from left to right in WT cPSE, WT cPAE, ACSS2^KO^ cPSE, and ACSS2^KO^ cPAE and the average thickness of DG lower blade region (right most) in males. **(D)** Neun+ staining in the hilar region of WT cPSE (upper left), ACSS2KO cPSE (upper right), WT cPAE (lower left), and ACSS2KO cPAE (lower right) and average abundance of neurons (right most) in males. Sample size n=4 biological replicates per sex per condition per genotype.

Interestingly, reduced hippocampal thickness in WT cPAE males was accompanied by an abnormal increase in the number of neurons in the hilus of the DG (**Figure 5D**), suggesting that potential impairments of neuronal migration and maturation might contribute to the observed reductions in thickness. This was only observed in WT males, while ACSS2^KO^ males exhibited a low number of hilar neurons irrespective of treatment (two-way ANOVA interaction: [genotype × treatment] F_1,12_=6.8, p=0.0244; post hoc Šidák: WT [cPSE vs cPAE] t_12_=3.138, adj. *p*=0.0086, cPSE [WT vs ACSS2^KO^] t_12_=3.554, p=0.0040, cPAE [WT vs ACSS2^KO^] t_12_=7.260, adj. *p*<0.0001) (**Figure 5D**). In females, there were no observable effects of treatment or genotype on hilus neuron counts **(Figure S5E).**

Overall, we found that cPAE significantly impacts hippocampal structure in both WT and ACSS2^KO^ mice, suggesting that alcohol affects hippocampal structure independently of the ACSS2 pathway.

### Loss of ACSS2 expression attenuates craniofacial abnormalities associated with cPAE

To capture macroscopic craniofacial phenotypes, WT and ACSS2^KO^ cPSE and cPAE mouse skulls were imaged using microCT and analyzed using 21 three-dimensional landmarks (**Figure S6A**). Euclidean Distance Matrix Analysis (EDMA) of linear distances revealed two significant differences in WT skulls following cPAE. The first, between landmarks 8 and 9, showed a 4.5% decrease of frontal bone width (95% Confidence intervals (CI) [0.904, 0.999], p<0.05) (**Figure 6A**), while the second indicated a 5.2% reduction in upper palate width between landmarks 12 and 13 (95% CI [0.904, 0.999], p<0.05) (**Figure 6B).** ACSS2^KO^ cPAE mice also displayed two significant local differences in skull development with cPAE: One between landmarks 5 and 16 showing a 4.5% increase in occipital bone length (95% CI [1.001, 1.086], p<0.05) and one between landmarks 2 and 3 indicating a 4% decrease in frontal bone length (95% CI [0.943, 0.978], p<0.05) (**Figures 6C,D**).

**Figure 6:**
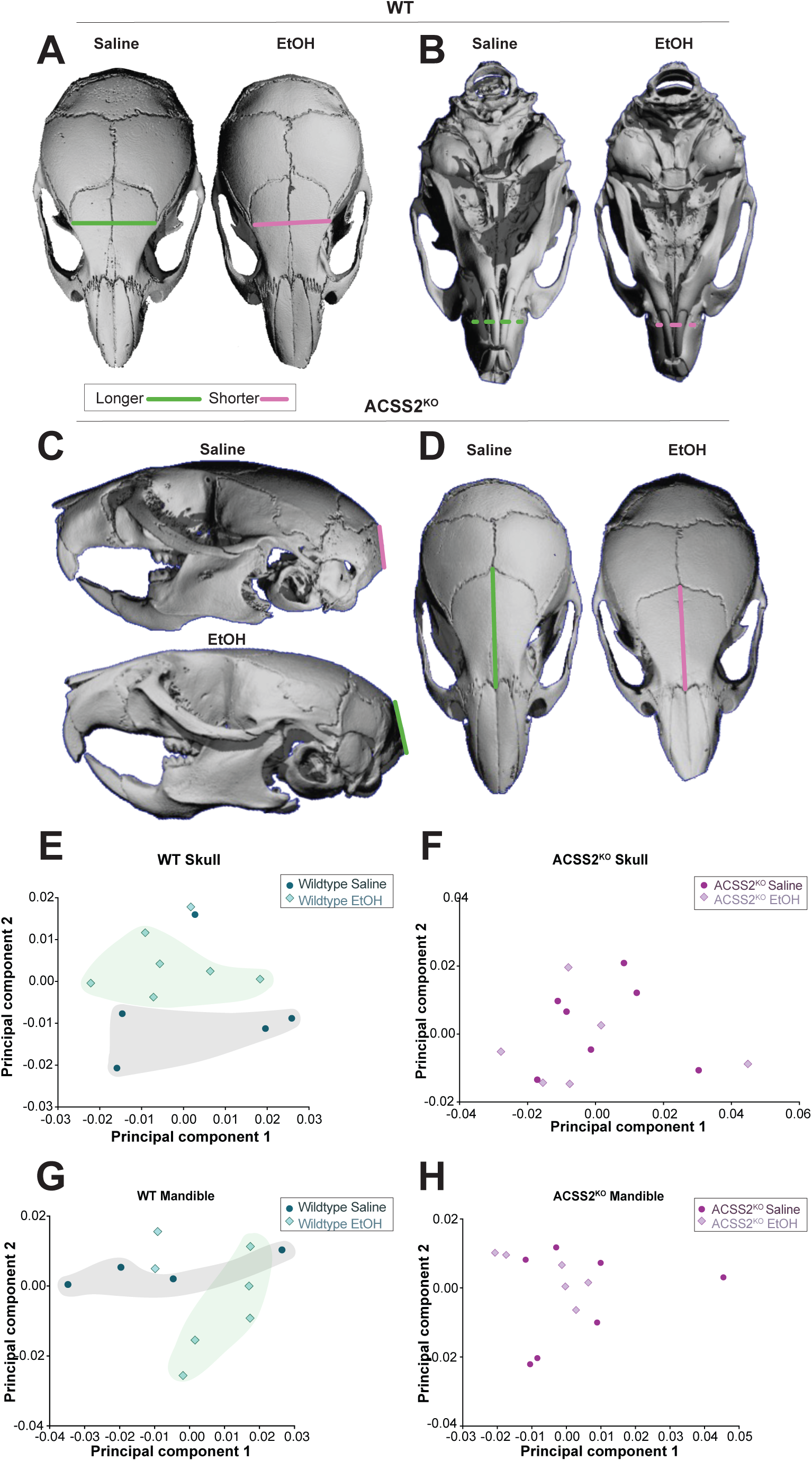
Loss of ACSS2 attenuates craniofacial variance associated with prenatal alcohol exposure. **(A-D)** Representative Micro-computed tomography (µCT) examples for significant Euclidean Distance Matrix Analysis (EDMA) results; green lines=significantly longer, pink lines=significantly shorter. **(A)** WT µCT representative examples of cPSE (left) and cPAE (right) and significant EDMA results across frontal bone landmarks 8 and 9. Sample size n=5 WT cPSE, n=7 WT cPAE **(B)** WT µCT images highlighting significant EDMA distances between landmarks 12 and 13. Sample size n=5 WT cPSE, n=7 WT cPAE **(C)** ACSS2^KO^ µCT representative examples of cPSE (upper) and cPAE (lower) and significant EDMA results along occipital bone landmarks 5 and 16. **(D)** ACSS2^KO^ µCT representative examples of cPSE (left) and cPAE (right) and significant EDMA results along frontal bone landmarks 2 and 3. Sample size n=7 ACSS2^KO^ cPSE, n=6 ACSS2^KO^ cPAE. **(E)** Principal component analysis (PCA) of WT cPSE and cPAE mice across principal component (PC) 1 and PC2, accounting for 54% of total variance across skulls. Sample size n=5 WT cPSE, n=7 WT cPAE. **(F)** PCA of ACSS2^KO^ cPSE and cPAE mice across PC1 and PC2, accounting for 70% of total variance across skulls. Sample size n=7 ACSS2^KO^ cPSE, n=6 ACSS2^KO^ cPAE. **(G)** PCA of WT cPSE and cPAE mice across principal PC1 and PC2, accounting for 72% of total variance across mandibles. Sample size n=4 WT cPSE, n=7 WT cPAE. **H)** PCA of ACSS2^KO^ cPSE and cPAE mice across PC1 and PC2, accounting for 73% of total variance across mandibles. Sample size n=7 ACSS2^KO^ cPSE, n=6 ACSS2^KO^ cPAE.

We also performed Principal Components Analysis (PCA) on landmark data to assess the general impact of cPAE on skull shape and found that WT mice displayed a larger spread of variance than their ACSS2^KO^ counterparts. In WT mice 35% of skull variance was captured by PC1 and 19% by PC2. We found a clear separation of cPAE and cPSE groups along PC2 (**Figure 6E**), indicating a strong overall effect of cPAE on craniofacial morphology. In contrast, there was no separation of ACSS2^KO^ mice by treatment along either PC1 (50% of variance) nor PC2 (20% of variance) (**Figure 6F**). We found a similar pattern when assessing differences of mandibular shape. In WT mice, 49% of variance was captured by PC1 and 23% by PC2, with a discernable separation of cPSE and cPAE mice along PC2 (**Figure 6G**). In ACSS2^KO^ mandibles 50% of variance was captured by PC1 and 23% by PC2, with no clear clustering by treatment along either PC (**Figure 6H**).

Taken together, these results indicate that craniofacial developmental changes associated with cPAE are ACSS2-dependent, suggesting an important role for alcohol-derived acetate in mediating these effects.

### Loss of ACSS2 mitigates cPAE-related memory and motor function deficits

To assess common motor and cognitive phenotypes associated with FASD, cPAE and cPSE mice underwent a battery of behavioral assays including open field, accelerating rotarod (AR), and Morris water maze (MWM). Open field showed no locomotor deficits in cPAE-treated WT and ACSS2^KO^ mice (**Figure S7A,B**), which is important for the interpretation of AR and MWM data.

The AR was used to assess differences in cerebellar motor function between groups, which has been previously shown to be affected by PAE.^40^ Repeated measures ANOVA revealed a sex × treatment interaction in WT mice (repeated measures ANOVA [treatment x sex] F_1,25_=4.363, *p*=0.047) (**Figure 7A**). While WT females showed no significant difference between cPAE and cPSE groups (**Figure 7B**), WT male cPSE mice had a longer latency to fall than cPAE counterparts across all trials (repeated measures ANOVA, male [cPSE vs cPAE] F_1,25_=6.292, *p*=0.019), largely driven by differences observed on day 1 (repeated measures ANOVA, WT Day 1 males [treatment] F_1,25_=7.502, *p*=0.044) and day 2 (repeated measures ANOVA, Day 2 WT males [treatment] F_1,25_= 7.68, *p*=0.044) (**Figure 7C**). In ACSS2^KO^ mice, we observed a slight reduction in the latency to fall in cPAE-treated mice on day 1 (Repeated measures ANOVA: ACSS2^KO^ Day 1 [treatment] F_1,16_=5.743, *p*=0.029) (**Figure 7D**). When stratifying by sex, ACSS2^KO^ female mice again showed no significant differences between cPSE and cPAE groups (**Figure 7E**). While cPAE ACSS2^KO^ males exhibited an initial motor deficit on day 1 of the AR (Repeated measures ANOVA: ACSS2^KO^ Males Day 1 [treatment] F_1,16_=26.942, *p*=8.9E-5), motor coordination was restored by day 2 **(Figure 7F),** suggesting that ACSS2 expression affects the development of cPAE-induced motor impairments in AR.

**Figure 7:**
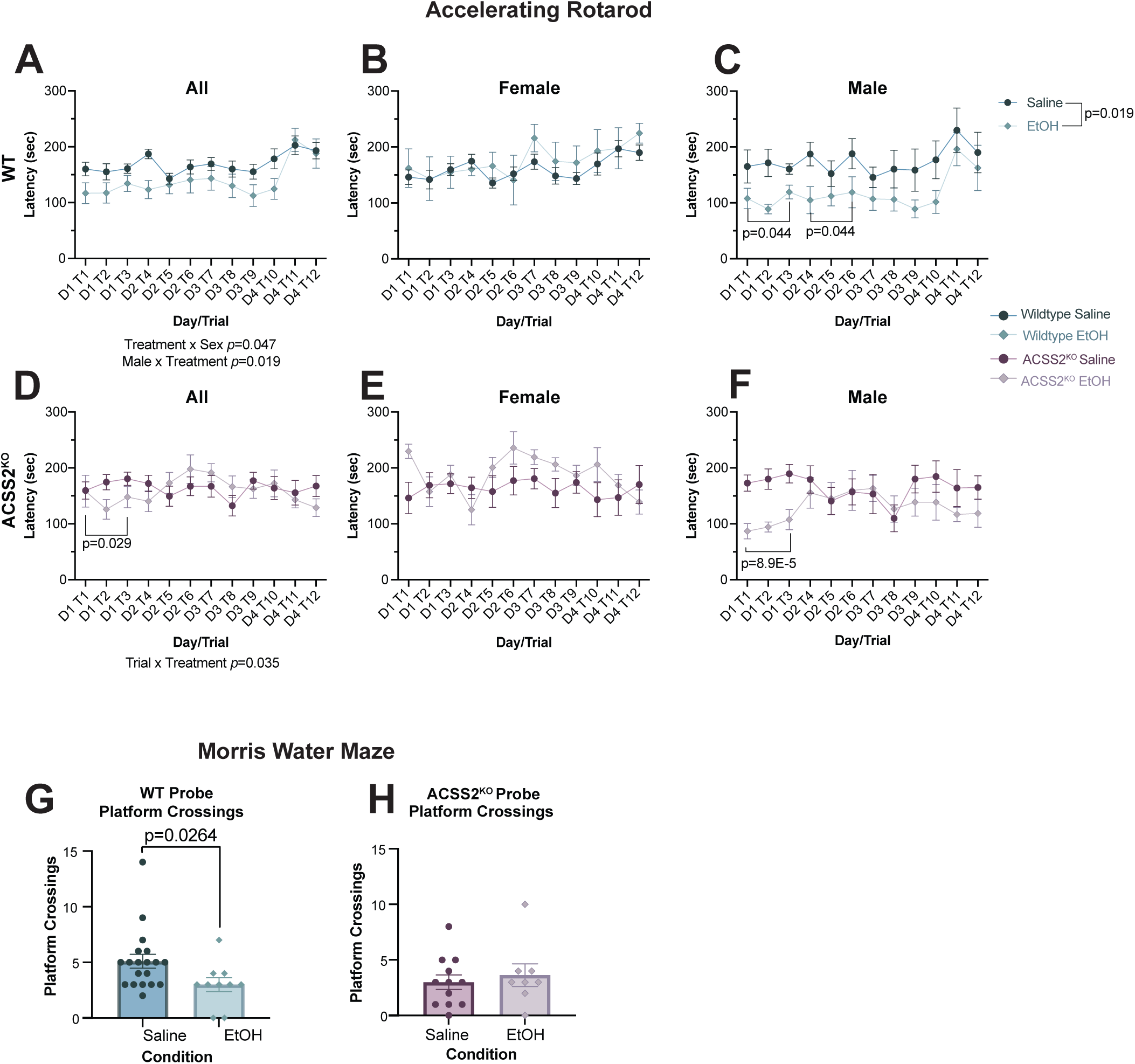
Loss of ACSS2 mitigates the effects of cPAE on motor function and spatial memory. **(A-F.)** Latency to fall of WT and ACSS2^KO^ mice following PSE or PAE during accelerated rotarod (AR) across four consecutive days (3 trials per day). **(A)** AR performance of all WT mice. Sample size n=19 cPSE and n=10 cPAE. **(B)** AR performance of female WT mice. Sample size n=13 cPSE and n=6 cPAE. **(C)** AR performance of male WT mice. Sample size n=6 cPSE and n=6 cPAE. **(D)** AR performance of all ACSS2^KO^ mice. Sample size n=12 cPSE and n=8 cPAE. **(E)** AR performance of female ACSS2^KO^ mice. Sample size n=6 cPSE and n=4 cPAE. **(F)** AR performance of male ACSS2^KO^ mice. Sample size n=6 cPSE and n=4 cPAE **(G-H)** Bar charts showing the average number of platform crossings in the Morris water maze probe trial between cPAE and cPSE groups in **(G)** WT mice (Sample size n=19 cPSE and n=10 cPAE), and **(H)** ACSS2^KO^ mice. Sample size n=12 cPSE, and n=8 cPAE.

The MWM was used to assess deficits of dorsal hippocampus-dependent spatial memory, which are hallmarks of FASD.^41,42^ We found that WT cPAE mice exhibited impaired long-term memory compared to cPSE counterparts, as evidenced by significantly fewer platform crossings during the probe trial (Wech’s t-test: [cPSE vs cPAE] t_23.82_= 2.367, *p*=0.0264) (**Figure 7G**). Further, WT cPAE mice exhibited an increased average distance from platform during the probe trial (Welch’s t-test: [cPSE vs cPAE] t_19,10_=2.151. *p*=0.0445) (**Figure S7C**). In contrast, we observed no effect of cPAE on either platform crossings (Welch’s t-test: [cPSE vs cPAE] t_12.58_= 0.5177, *p*=0.6317) (**Figure 7H**) nor average distance from platform (Welch’s t-test: [cPSE vs cPAE] t_18,9_=1.085. *p*=0.8926) (Figure S7D) in ACSS2^KO^ mice. cPAE did not affect swim speed in either genotype (**Figure S7E,F**). Thus, our results suggest that cPAE-induced impairments of dorsal hippocampal long-term memory depend on ACSS2.

Taken together, we found that loss of ACSS2 mitigates behavioral impairments related to cPAE, suggesting that ACSS2-mediated metabolic-epigenetic effects play an important role in motor and memory deficits associated with FASD.

## DISCUSSION

Here, we establish the functional importance of an emerging metabolic-epigenetic pathway in key phenotypes related to PAE. We identify the developmental window for the nuclear translocation of ACSS2 and show that alcohol-derived acetate is deposited on histones in the developing brains. Using a genetically modified ACSS2^KO^ mouse, we show that this pathway is required for persistent transcriptional alterations and the manifestation of FASD hallmarks including abnormal skull development, reduced motor function, and spatial memory deficits. Together, our results outline a novel aspect of FASD etiology and highlight the previously unknown role of ACSS2 and alcohol-derived acetate in the development of these disorders.

Our findings align with earlier reports on the roles of alcohol metabolism and epigenetics in FASD. Metabolic reactions that convert ethanol into acetaldehyde and subsequently acetate are accompanied by the generation of reactive oxygen species (ROS) which have been tightly linked to DNA damage.^43^ Thus, ROS are thought to play an important role in neurodevelopmental impairments related to FASD.^44^ Treatment of FASD in rodents with antioxidants such as Vitamin E and C reduces ROS and alleviates certain symptoms including brain and liver atrophy.^45,46^ Further, PAE was shown to affect epigenetic mechanisms, in particular histone acetylation.^47–49^ In addition to ROS production,^50^ prenatal alcohol-induced epigenetic changes were previously attributed to altered availability of substrates and cofactors ^51–53^ and increases in the NADH/NAD ratio. ^52,54^

We now show that ACSS2 plays a critical role in regulating core FASD phenotypes and related epigenetic and transcriptional changes. This is in line with the recently emerging role of this pathway in alcohol use-related behaviors during postnatal alcohol exposure. Specifically, ACSS2-mediated incorporation of alcohol-derived acetate into brain histone acetylation is required for the formation of alcohol-associated memories^17^ and genetic deletion of ACSS2 reduces voluntary alcohol intake in mice.^19^ We now demonstrate that loss of ACSS2 prevents alcohol-related worsening of memory in mice prenatally exposed to alcohol. These results are remarkable as ACSS2 and acetate were shown to promote^19^ memory formation in adult mice in other contexts.^24,55,28^ Indeed, overexpression of ACSS2^56^ or dietary supplementation of exogenous acetate^57,56^ both rescue memory in mouse models of Alzheimer’s disease, while inhibition of this pathway has been linked to worsened molecular and behavioral outcomes.^57,56^ In stark contrast, loss of ACSS2 attenuates epigenetic and transcriptional changes and is beneficial for memory in the context of PAE, suggesting that the same metabolic-epigenetic pathways that usually promote memory could be detrimental following exposure to chronic metabolic stress developmentally.

Intriguingly, we found that loss of ACSS2 did not uniformly impact all phenotypic hallmarks of FASD. In addition to its effect on memory, ACSS2 KO also reduced motor function impairments and the formation of craniofacial abnormalities in cPAE mice. Previously, ACSS2 has been largely studied in the context of hippocampal memory, and thus our current study is the first to link this pathway to implicit motor learning regulated by the cerebellar vermis.^58,59^ The cerebellum is strongly impacted by alcohol both postnatally^60^ and prenatally^61^ and our findings indicate that these impairments are in part mediated by ACSS2. Intriguingly, the beneficial effect of ACSS2 KO on motor function was specific to males, which is in line with emerging reports of differential metabolic-epigenetic regulation between the sexes^19,28,62,63^ and could in part contribute to previously reported sex-specific manifestations of FASD.^64,65^

Craniofacial phenotypes characteristic of FASD^4^ are largely thought to be the result of improper neural crest cell migration and apoptosis.^66^ The contribution of alcohol-related metabolic-epigenetic interactions is entirely novel, revealing an unexpected mechanism underlying FASD. Intriguingly, other PAE-related phenotypes such as reduced hippocampal thickness were not affected by ACSS2 deletion. This suggests that several alcohol-related molecular pathways^67^ act in parallel to orchestrate the development of FASD, and their relative contribution to specific phenotypes warrants further investigation.

ACSS2 has been previously linked to differentiation of neuron-like cells,^24^ its role in more physiological developmental contexts, however, remains unknown. Here, we show that ACSS2 translocates into nuclei around E11.5 in the developing brain, which coincides with peak neurogenesis.^21,22^ To our knowledge, our study represents the first identification of a developmental window for the nuclear translocation of a metabolic enzyme. Importantly, while nuclear translocation of metabolic enzymes has been described in several cell types, tissues and organisms,^13–16,68^ the necessity of this re-localization for their epigenetic impact has been contested in the field. Here, we find that alcohol-derived acetate can be incorporated into brain histone acetylation prior to the nuclear and chromatin localization of ACSS2, suggesting that cytoplasmic expression of this enzyme might be sufficient to mediate histone acetylation. This is contrary to previous reports suggesting a clear segregation between nuclear and cytoplasmic acetyl-CoA pools^69^ and decreased histone H3 acetylation following nuclear exclusion of ACSS2.^70^ Overall, these seemingly contradictory findings highlight the need for a better mechanistic understanding of how intermediary metabolites generated by ACSS2 and other metabolic enzymes influence epigenetic processes.

We found that cPAE impacted the epigenome in both the dHPC and the vermis in an ACSS2-dependent manner. The most significantly affected hippocampal hPTM was H2A.Z acetylation, which recently emerged as a key regulator of learning and memory^71–73^. cPAE-induced H2A.Z acetylation was dysregulated in both brain regions, and, in line with the observed behavioral outcomes, this effect was blunted in ACSS2^KO^ mice. Importantly, ACSS2-dependent acetylation of the histone variant H2A.Z was previously unknown, as this enzyme has primarily been linked to histone H3 and histone H4 acetylation.^17,24,74^ Metabolic regulation of H2A.Z acetylation is further supported by our recent findings outlining a critical role for this histone variant in acetate-enhanced memory.^28^

Long-lasting epigenetic remodeling that persists into adulthood could be a key mechanism underlying FASD. We found that cPAE resulted in decreased enrichment of several histone H3 and H4 acetylation marks in the hippocampus. In contrast, acute alcohol exposure generally increases histone H3 and H4.^75^ This difference is likely due to compensatory mechanisms driven by repeated long-term exposure to ethanol, as well as epigenetic differences between the developmental stages when alcohol is administered. Indeed, it has been previously shown that acute and chronic exposure to heroin results in opposing histone acetylation changes in the striatum of rats.^76^ Of note, histone acetylation was largely unaffected in the ACSS2^KO^ brain in our study, again underscoring the role of metabolic-epigenetic interactions in cPAE-related chromatin changes. In further support, both histone acetyltransferase and histone deacetylase levels were unchanged in the hippocampus of cPAE mice in both our and other studies,^77^ suggesting that altered epigenetic modifiers are unlikely to be responsible for the observed chromatin modifications.

Prenatal exposure to alcohol results in lasting transcriptional changes, which we also observed in our model. Interestingly, we found limited enrichment of GO terms across gene and transcript analyses. This is in line with previous studies reporting a large number of DEGs resulting in few functional themes across repeated exposure paradigms^78,79^ suggesting broad effects on transcription impacting many independent pathways. Importantly, loss of ACSS2 had a profound effect on PAE-related gene expression changes, which aligns with the emerging role of this metabolic enzyme as a key transcriptional regulator.^17,24,80^ We found a consistently fewer DEGs and DETs in ACSS2^KO^ mice compared to WT, in line with the attenuated behavioral and morphological impairments. Strikingly, few DEGs and DETs overlapped between genotypes and all but one differed in directionality. Further, we identified hundreds of DEGs and DETs exhibiting significant treatment ^x^ genotype interactions. Of note, deletion of ACSS2 in AUD paradigms generally results in increased number of affected DEGs,^19,24^ suggesting that the role of this pathway in mediated ethanol-induced gene expression changes depends on the developmental stage. Further, our study is one of the first to investigate the effect of cPAE on alternative splicing, highlighting an additional 361 genes. Alternative splicing events and transcription in the context of PAE have recently emerged,^81^ however, our study is the first of its kind to investigate the contributions of alternative splicing by comparing DEGs and DETs following cPAE.

Taken together, our results outline a previously unknown aspect of FASD etiology driven by ACSS2. We show that this pathway regulates core phenotypes, warranting further investigation of metabolic-epigenetic interactions in the context of PAE. Emphasizing the key role of metabolism, the essential nutrient choline has been tightly linked to neurodevelopment, especially in the hippocampal cholinergic system, and choline supplementation enhances dendritic arborization, neuronal function and cognition.^82,83^ Indeed, choline supplementation was shown to have cognitive benefits in FASD patients.^84^ By linking metabolism to epigenetic processes in the context of PAE and by demonstrating the functional importance of these emerging pathways, our study outlines a novel aspect of FASD biology that will inform the development of future therapies.

## ACKNOWLEDGEMENTS

MicroCT imaging was performed at the Musculoskeletal Research Center at Washington University in St. Louis; we thank Michael Brodt for technical assistance. Behavioral research reported in this manuscript was performed in the Animal Behavioral Subunit of the Intellectual Developmental Disabilities Research Center at Washington University in St. Louis (IDDRC@WUSTL, NICHD P50HD103525); we thank Katie McCullough for technical assistance. RNA sequencing was performed at the McDonnell Genome Institute at Washington University in St. Louis. This work was supported by NIH grants R00AA028577 (GE), Alzheimer’s Association grant AARF-19-618159 (GE), 7R01AI118891 (BAG), R01HD106051 (BAG), IDDRC@WUSTL Flagship Pilot Project Grant (GE), the NARSAD Young Investigator Grant YIG31527 from the Brain and Behavior Research Foundation (GE).

## Author contributions

KMD, conceptualization, investigation, visualization, formal analysis and writing; EMP, investigation, visualization, formal analysis, writing; AY, ML, AJB, AO, CC and FNV, investigation, visualization, formal analysis; BAG, funding acquisition, supervision; CH and SM, conceptualization, supervision, formal analysis; GE, conceptualization, investigation, visualization, formal analysis, funding acquisition, supervision and writing.

## STAR METHODS

## KEY RESOURCES TABLE

0 KEY RESOURCES TABLE
| Resource | Source | Identifier |
| --- | --- | --- |
| Antibodies |  |  |
| ACSS2 Monoclonal Antibody | Invitrogen | Cat. No. MA5-14810, Lot No. ZL4583521 |
| Goat anti-Guinea Pig IgG (H+L) Highly Cross-Adsorbed Secondary Antibody, Alexa Fluor™ 594 | ThermoFisher Scientific | Cat. No. A-11076 |
| Goat anti-Rabbit IgG (Heavy chain) Superclonal™ Secondary Antibody, HRP | Invitrogen | Cat. No. A27036, Lot No. 2961376 |
| NeuN antibody | Synaptic Systems | Cat. No. 266 004 |
| Biological Samples |  |  |
| C57BL/6J fetal brain tissue E10.5, E11.5, E17.5 | This paper | N/A |
| C57BL/6J adolescent brain tissue | This paper | N/A |
| C57BL/6J ACSS2<sup>KO</sup> adolescent brain tissue | This paper | N/A |
| Chemicals, peptides, and recombinant proteins |  |  |
| AEBSF | Sigma | Cat. No. A8456-25MG |
| Microcystin | ThermoFisher Scientific | Cat. No. 502008727 |
| NP-40 (IGEPAL CA-630) | Sigma | Cat. No. I8896 |
| SeeBlue Plus 2 Ladder | ThermoFisher Scientific | Cat. No. LC5925 |
| Sequencing Grade Modified Trypsin | Promega | Cat. No. V5113 |
| Propionic Anhydride | Sigma | Cat. No. 800608 |
| 15-well 4-12% Bis-Tris Gel | ThermoFisher Scientific | Cat. No. NP0323BOX |
| Ponceau Solution | Sigma | Cat. No. P7170-1L |
| 1M Sodium Butyrate | Sigma | Cat. No. B5887-250MG |
| TRIzol | Invitrogen | Cat. No. 15596026 |
| Chloroform | Sigma | Cat. No. C2432-25mL |
| Qiagen RNase-free DNase | Qiagen | Cat. No. 79254 |
| Normal Goat Serum (NGS) | Abcam | Cat. No. ab7481 |
| Ethylene-glycol | ThermoFisher Scientific | Cat. No. A11591.36 |
| Critical commercial assays |  |  |
| Qiagen RNeasy Miniprep Kit | Qiagen | Cat. No. 74106 |
| Qubit Protein Assay Kit | ThermoFisher Scientific | Cat. No. 74106 |
| Subcellular Fractionation Kit for Tissues | ThermoFisher Scientific | Cat. No. Q33211 |
| NuPAGE SDS Buffer Kit | ThermoFisher Scientific | Cat. No. 87790 |
| SuperSignal™ West Pico PLUS Chemiluminescent Substrate | ThermoFisher Scientific | Cat. No. NP00060 |
| NEBNext® Poly(A) mRNA Magnetic Isolation Module | New England Biolabs | Cat. No. E7490 |
| NEBNext® Ultra™ II RNA Library Prep Kit for Illumina® | New England Biolabs | Cat. No. E7770 |
| D1000 DNA ScreenTape Analysis | Agilent | Cat. No. 5067-5602 and 5067-5582 |
| NEBNext® Library Quant Kit | New England Biolabs | Cat. No. E7630 |
| HighPrep PCR beads | MagBio Genomics | Cat. No. AC-60050 |
| Deposited Data |  |  |
| Raw data files for LC-MS/MS targeting histone peptides | This paper | MassIVE |
| RNA sequencing data | This paper | GEO |
| Experimental models: Organisms/Strains |  |  |
| Wildtype, C57/BL6J | The Jackson Laboratory | 000664 |
| ACSS2<sup>KO</sup>, C57BL6J | Alexander et al., 2022 | 000664 |
| Oligonucleotides |  |  |
| NEBNext® Oligos for Illumina | New England Biolabs | Cat. No. E7335 and E7500 |
| Software and Algorithms |  |  |
| Adobe Illustrator | Adobe creative cloud | N/A |
| Any Maze | Stoelting Co. | Item no. 60000 |
| Microsoft Office Suite | Microsoft | n/a |
| FreezeFrame | Actimetrics | N/A |
| GraphPad Prism v10.50 | GraphPad Prism | N/A |
| LAS X Imaging Software | Leica Microsystems | N/A |
| MorphoJ | Klingenberg, 2011 | N/A |
| R package: ComplexHeatmap | Gu, 2022 | N/A |
| R package: DESeq2 | Love, Huber, and Anders, 2014 | N/A |
| R package: ggplot2 | Wickham et al., 2016 | N/A |
| R package: rrvgo | Sayols, 2023 | N/A |
| R package: topGO | Alexa and Rahnenfuhrer, 2024 | N/A |
| SPSS Statistics | IBM | N/A |
| featureCounts 2.0.6 | Liao, Smyth, and Shi, 2014 | N/A |
| stringApp in Cytoscape v3.10.3 | Doncheva et al., 2019 | N/A |
| 3D Slicer | Slicer.org | N/A |
| Motor Monitor | Kinder Scientific | N/A |
| Other |  |  |
| AccuRotor EzRod | Omnitech Electronics Inc. | N/A |
| GRCm39 RefSeq assembly | NCBI | Assembly accession GCF_000001635.27 |

## RESOURCE AVAILABILITY

### Lead contact

Further information and requests for resources and reagents should be directed and will be fulfilled by the Lead Contact Dr. Gabor Egervari

### Materials availability

This study did not generate new or unique reagents or materials.

### Data and code availability

All proteomic and genomic data will be made publicly available at time of publication.

## EXPERIMENTAL MODEL

### Animal Models

Female mice were housed with studs for five nights. Following night one, female mice were injected intraperitoneally with 3g/kg EtOH from a 20% EtOH solution or an equivalent volume of 1x PBS until they gave birth. Once visibly pregnant, dams were isolation housed until pup weaning at postnatal day (PND) 21. Offspring were euthanized between PND40-50 via cervical dislocation or transcardial perfusion. Following cervical dislocation brains were extracted and flash-frozen in 2-methyl butane before being stored at -80°C. Following fixation, heads were removed and placed in 4% PFA for one week at 4°C before being placed in 1x PBS at 4°C until MicroCT imaging.

Fetal brain samples were collected from timed pregnancies at embryonic days (E) 10.5, 11.5, and 17.5. Dams were injected intraperitoneally with 4g/kg of 20% EtOH in 1x PBS, 4g/kg of 20% D6-EtOH in 1x PBS, or an equivalent volume of 1x PBS four hours prior to euthanasia. Dams were sacrificed via cervical dislocation; brains were extracted and flash frozen in 2-methyl butane. Using fine tip forceps (FST, Cat. No.11412-11) embryos were removed; E10.5 and E11.5 brains were grouped into pools of whole fetal brain, while E17.5 brains were dissected into pools of forebrain and midbrain. All tissues were flash-frozen in 2-methyl butane and stored at -80°C.

## METHOD DETAILS

### Histone Extraction

Histones were extracted following the previously described protocol^85^ with the following alterations: Samples were homogenized in 200µL of Nuclear Isolation Buffer (NIB) (15 mM Tris-HCl, 15 mM sodium chloride, 60 mM potassium chloride, 5 mM magnesium chloride, 1 mM calcium chloride, and 250 mM sucrose at pH 7.5; with 0.5 mM AEBSF, 10 mM sodium butyrate, 5 mM microcystin, and 1 mM dithiothreitol) and 10% NP-40.

Following this, homogenates were centrifuged (600rcf for 5min at 4°C) and the supernatant was discarded. Nuclear pellets were resuspended in 200µL NIB and centrifugation repeated. Pellets were resuspended in 200µL of 0.4N sulfuric acid and rotated at 4°C for 4 hours. Samples were centrifuged (3,400rcf for 5 min at 4°C) and supernatant was collected. Histones were precipitated by adding 50µL of 100% Trichloroacetic acid and incubating samples in ice at 4°C overnight. Samples were washed with 0.1% HCl in acetone, centrifuged (3,400rcf for 5 min at 4°C), supernatant discarded, and repeated with 100% acetone (4,000rcf for 2 minutes). Acetone was removed and histones were air-dried for 20 before resuspension in30µL of ammonium bicarbonate. Chemical derivatization, trypsin digestion, propionylation of N-termini, desalting, and analysis of histone peptide steps were performed as previously described.

### Subcellular Fractionation

Subcellular fractionation was performed using Subcellular Fractionation Kit for tissues (ThermoFisher, Cat. No.87790) following manufacturer protocols with the following modifications: buffer volumes were proportioned for 10mg of protein (one fetal brain pool); after cytoplasmic extraction all steps were completed in a 1.5mL microcentrifuge tube. To remove excess lipids, cytoplasmic extracts were precipitated in 200uL of acetone at - 20°C for one hour. Samples were centrifuged (14,000 rcf for 10 minutes), and supernatant discarded. Pellets air-dried for 20 minutes and were resuspended in 65µL of molecular grade water. All fractions were stored at - 80°C.

### Western Blot

Protein was quantified with the Qubit Protein Assay (ThermoFisher, Cat. No. Q33211) and Qubit Fluorometer 4, according to manufacturer protocol. Samples were prepared using NuPAGE SDS Buffer Kit (ThermoFisher, Cat. No.NP00060) with 8µg of protein, 3.75µL of 4x LDS buffer, 1.5µL sample reducing agent, and molecular grade water up to 15µL. Samples were denatured at 95°C for five minutes. The gel (ThermoFisher, Cat. No. NP0323BOX) was loaded into XCell minilock (ThermoFisher, Cat. No. EI0001) and filled with 1x MOPS running buffer. 6µL of SeeBlue Plus 2 Ladder (ThermoFisher, Cat. No. LC5925) was added as a standard; samples were loaded into the remaining lanes before running electrophoresis for 2.5 hours at 100V. The gel was wet transferred onto nitrocellulose membrane (Bio-Rad, Cat. No. 1620112) at 25V for 90 minutes. Following transfer, membrane was stained with Ponceau and imaged with the iBright™ FL1500 Imaging System (Invitrogen, Cat. No. A44241). The blot was rinsed with 1xTBST and blocked (5% Milk 1x TBST) for 1 hour at RT, shaking. The membrane was agitated overnight at 4C in 1:1000 ACSS2 (Cat. No MA5-14810, Lot No. ZL4583521) in 5% Milk 1x TBST solution. The membrane was washed 3x in 1x TBST for 10 minutes each, before being placed in 1:5000 Goat anti-Rabbit IgG (Heavy chain) Superclonal™ Secondary Antibody, HRP (Cat. No A27036, Lot No. 2961376) in 5% Milk 1x TBST Solution. The membrane was washed in TBST 3x, 10 minutes each before SuperSignal™ West Pico PLUS Chemiluminescent Substrate (Cat No. 34580) was applied according to manufacturer protocols. The membrane incubated at RT for 5 min and was then imaged on the iBright™ FL1500 Imaging System. Blots were quantified relative to their total histone volume in the chromatin-bound fraction of the Ponceau stained blot. Analysis of bands was completed via Excel and GraphPad Prism.

### RNA Extraction and Sequencing

Total RNA was extracted using TRIzol-chloroform and the RNeasy Mini Kit (Qiagen, 74104) with RNase-free DNase (Qiagen, 79254) treatment according to manufacturer protocols. Total RNA quality was assessed on an Agilent 4200 TapeStation using the High Sensitivity RNA ScreenTape Assay (Agilent, 5067-5579 and 5067-5580).

mRNA was isolated from 10 ng total RNA using the NEBNext Poly(A) mRNA Magnetic Isolation Module (New England Biolabs, E7490), and libraries were prepared using the NEBNext Ultra II Directional RNA Library Prep Kit for Illumina (New England Biolabs, E7760) in coordination with NEBNext Multiplex Oligos for Illumina (New England Biolabs, E6440) and HighPrep PCR Beads (MagBio, AC-60050). Library quality was assessed using the D1000 ScreenTape Assay (Agilent, 5067-5582). Libraries were quantified using the NEBNext Library Quant Kit (New England Biolabs, E7630) on an Applied Biosystems QuantStudio 5 Real-Time PCR System. The concentration of amplifiable templates in each library was calculated using the openly available NEBioCalculator qPCR Library Quantification Web Tool. Following quantification, libraries were pooled in equal concentrations and submitted for sequencing.

Libraries were sequenced on an Illumina NovaSeq X Plus using paired end reads extending 150 bases, and base calls and demultiplexing were performed with Illumina’s bcl2fastq software with a maximum of 1 mismatch in the indexing read at the Genome Technology Access Center (GTAC) at the McDonnell Genome Institute (MGI) at Washington University School of Medicine in St. Louis.

### Immunohistochemistry

Following MicroCT imaging, skulls were removed, and brains were stored in 30% sucrose in 1x PBS for at least 48 hours. Brains were sectioned to 25µm thickness with the Leica CM1950 Cryostat at -20°C. Slices were deposited into a 12-well plate containing 2mL cryoprotectant (28% sucrose, 30% Ethylene glycol in saline) using a fine paint brush. Plates were covered in foil and stored at -20°C. Brain slices were washed 1x PBS for 10 minutes, then 3x in 5%Tween20-1xPBS for 15 minutes at room temperature. Sections were shaken for an hour in 500µL blocking solution (10% Normal Goat Serum (NGS) (Abcam, Cat. No. ab7481), 0.3M Glycine in 5% Tween20-1xPBS. Blocking solution was removed and sections stained with 1:500 NeuN (Synaptic Systems, Cat. No. 266 004) in 10mM glycine in 0.5% Tween20-1xPBS solution shaking overnight at 4°C. Washing and blocking were repeated. Slices were transferred to new wells and incubated in 500µL of 1:1000 goat anti-guinea pig 594 (ThermoFisher, Cat. No. A-11076) in 2% NGS in PBST-Tween20. Slices were wet mounted, coated with DAPI mounting media (Southern Biotech, Cat. No. 0100-20) and sealed. All slices were imaged using the Leica DMi8 inverted microscope paired with LAS X imaging software v5.3.1 at 20X magnification 12x4 tiles, and a 25µM Z-stack. Images were blindly analyzed for hippocampal thickness and NeuN+ counts using ImageJ. Data visualization, and analysis were completed using GraphPad Prism and Adobe Illustrator.

### MicroCT

Whole skulls were scanned on the SCANCO µCT50 for 2.1 hours each at room temperature, before being placed back in 1xPBS at 4°C. Whole mouse heads were imaged at the Musculoskeletal Research Center at Washington University in St. Louis using the SCANCO µCT50. Scans were exported as DICOM files and analyzed using 3D Slicer. 3D renderings of mouse skulls were completed using the evaluate and ray tracer SCANCO µCT50 software according to manufacturer protocols.

### Behavioral Tasks

All behavioral testing was conducted in the IDDRC Animal Behavior Subunit at WashU Medicine. Behavior was assessed in two independent cohorts of mice. Cohort 1 consisted of 29 mice, 10 EtOH-exposed (4F, 6M) and 19 PBS-exposed (13F, 6M). Cohort 2 consisted of 20 mice, 8 EtOH-exposed (4F, 4M) and 12 PBS-exposed (6F, 6M). Mice were handled for 3-5 days prior to the behavioral testing and the tails of mice were marked with a non-toxic, permanent marker during weight collection and regularly thereafter to easily distinguish mice during testing. For all tasks, the mice were acclimated to the testing room at least 30 minutes prior to the start of testing. Testing orders were randomly counterbalanced for genotype across apparatuses and trials. All mice were behaviorally tested as juveniles starting at P34-36. All assays were conducted by a male experimenter blinded to experimental group designations, and all testing occurred during the light/inactive phase. Tasks were ordered from least to most stressful to minimize the order-of-testing effects: 1-hr locomotor activity, accelerating rotarod, water maze, fear conditioning. Unless otherwise indicated, all equipment was cleaned between animals with a 0.02% chlorhexidine diacetate solution (Nolvasan, Zoetis) or 70% ethanol solution.

#### 1-hour Locomotor Activity and Exploration

Activity levels were assessed for a single, hour-long trial based on our published methods^1^. Briefly, horizontal and vertical beam breaks were quantified as ambulations and rearings, respectively, by computer software (MotorMonitor, Hamilton-Kinder, LLC, Poway, CA) in a 33 × 11 cm central zone and a bordering 5.5 cm peripheral zone. General activity variables (total ambulations, rearings, time at rest), along with measures of emotionality, including time spent, distance traveled, and entries made into the central zone were analyzed.

#### Accelerating Rotarod

Cerebellar-based motor coordination and motor learning were assessed using the accelerating rotarod, based on previous methods.^86,87^ The AccuRotor EzRod apparatus (Omnitech Electronics, Inc) was used. Animals received three trials a day on each of four consecutive days on an accelerating rod (4-40 rpms) for 300 seconds. An intertrial interval of 30 minutes was used each day. Each animal received a habituation trial on a continuously rotating rod at 3rpms for 60 seconds prior to the first test trial on the first day only. The duration the animal could remain on the rod was measured by Fusion software (Omnitech Electronics, Inc).

#### Morris Water Maze

Spatial learning and memory were assessed using Morris Water Maze (MWM), based on previously published methods.^88^ Briefly, Cued, Place, and Probe trials were conducted in a white plastic pool measuring 124 cm in diameter (polyethylene; DuraCast manufacturer). White tempera paint was used to opaque the water. The PVC escape platform measured 11.5 cm in diameter. An overhead digital video camera connected to a PC computer, and the computer software program ANY-maze (Stoelting Co., Wood Dale, IL) tracked the swimming pathway of the mouse to the escape platform and quantified path length, latency to find escape platform, and swimming speeds. Cued trials were conducted over two consecutive days and consisted of 4 trials separated by 30 min intervals. Cued trials serve to acclimate the mice to the pool, confirm swimming ability and vision. A red tennis ball on a pole was attached to the center of the escape platform and served as a visual cue. The platform was moved to a different quadrant for each trial to prevent spatial acquisition. The mouse was dropped off from the quadrant opposite the escape platform in each trial, was given 60 s to find the platform and 10 s on the platform before being placed into warming cages. Three days after, Place trials were conducted daily for five days consisting of two blocks of two consecutive trials with an hour in between blocks. The escape platform stayed in the same location over all trials and distal cues were placed on the surrounding walls to assist in spatial learning. The drop off location was different for each of the four trials per day. Trials were 60 s with 10 s on the platform to aid spatial learning, before being placed in warming cages. An hour following the completion of the last Place trial, a single 60 s Probe trial assessed memory retention of the escape platform, which was removed but distal cues remained.

## QUANTIFICATION AND STATISICAL ANALYSIS

### Histone LC-MS/MS

To detect heavy labeling raw MS files were analyzed manually at the three most reliably detected acetylated peptides (H3K14ac, H3K23ac, and H4K16ac). Each mark was further analyzed using a two-way ANOVA (timepoint vs condition) and in the case of significant (*p*<0.05) interaction, pairwise differences were explored using post hoc Šídák’s multiple comparisons testing. Statistical analysis of histone ratios of LC-MS/MS-derived data was completed using Welch’s t-test for relative abundance of histone peptide ratios comparing saline and EtOH groups per genotype per region. Individual hPTMs were further explored using two-way ANOVA (genotype vs condition) followed by post hoc Šídák’s multiple comparisons test to investigate pairwise differences in the case of significant (*p*<0.05) interaction. Data visualization and curation was completed using Excel, GraphPad Prism, and Adobe Illustrator.

### RNA-seq

Adapters were trimmed from reads using Cutadapt 5.0 with Python 3.12.10^89^ with a minimum length of 15 bp. Reads were aligned to the GRCm39 *Mus musculus* reference genome assembly using STAR 2.7.10a^90^ Counts at gene and transcript levels were quantified using featureCounts 2.0.6^91^ and the annotation from the NCBI RefSeq assembly GCF_000001635.27. Before differential expression analysis (DEA), counts were filtered to remove rows with low counts, keeping only rows meeting criteria of at least 2 samples within a genotype having at least 5 read counts each.

DEA for gene and transcript levels was performed with DESeq2^92^ using the default Wald negative binomial test to determine differentially expressed genes (DEGs) or differentially expressed transcripts (DETs) for pairwise comparisons (cPAE vs. cPSE within a genotype and region) and genotype × exposure interaction (within each region). Unless otherwise stated, an adjusted p-value cutoff of 0.1 and no log_2_(fold change) cutoff was used in distinguishing differentially expressed from non-differing features to maximize the sensitivity of RNA-seq results. Unless otherwise specified, the R package ggplot2^93^ was used to generate plots of RNA-seq data.

Gene ontology (GO) enrichment analysis was performed using topGO^94^ with biological process (BP) ontology, minimum node size of 10, weight01 algorithm, and Fisher’s exact test. Raw p-values were adjusted using the Benjamini-Hochberg method, and terms with FDR<0.1 were considered significantly enriched. When >10 significant terms were identified, redundancy was reduced using the R package rrvgo^95^ as follows: semantic similarity between terms was calculated, a similarity matrix was generated (Lin method), terms were clustered (similarity threshold 0.7), and representative nonredundant terms (one parent term per cluster) were retained.

For heatmaps, counts were normalized using blind variance stabilizing transformation (VST) generated by DESeq2. Heatmaps were generated using ComplexHeatmap^96^ and show Z-score of VST-normalized group average counts.

To generate protein-protein interaction (PPI) networks, STRING protein query in Cytoscape was used with the following parameters: Mus musculus, full STRING network, confidence (score) cutoff 0.4, maximum additional interactors. Network diagrams were generated using stringApp in Cytoscape v3.10.3.^97^ RNA-seq results were imported into Cytoscape to map node attributes. Network layouts were set using prefuse force directed OpenCL layout based on STRING neighborhood score, and edge thickness was mapped to STRING composite score. Only clusters containing ≥5 nodes are shown.

Differential alternative splicing events (ASEs) were identified between cPAE and cPSE using rMATS v4.3.0^98^ with variable read lengths and soft-clipped reads permitted. Change in percent spliced in (ΔPSI) was

### Immunohistochemistry

Statistical analysis of neuron quantity and hippocampal thickness were analyzed using a two-way ANOVA (genotype × condition) stratified by sex. Post hoc Šídák’s test was used to investigate pairwise differences in the case of significant (*p*<0.05) interaction. Data visualization and curation was completed using GraphPad Prism and Adobe Illustrator.

### MicroCT PCA and Anatomical Configuration

A total of 25 mice at 45 days of age were fixed and whole mouse heads were scanned at the Musculoskeletal Research Center at Washington University in St. Louis. Twenty-one landmarks (16 cranial, five mandibular) were collected from three-dimensional reconstructions of the cranium and left hemi-mandible using 3D Slicer following previously established methods.^99,100^ Mandibular and craniofacial landmarks were also analyzed separately. Skulls from the wildtype group and the ACSS2^KO^ mice were compared to identify patterns of differences in the mice. Principal components were calculated from a generalized Procrustes analysis using MorphoJ software.^101^ Euclidean Distance Matrix Analysis was completed to identify differences in individual linear distances.

### Behavior

Behavioral statistical analyses and data visualization for data were conducted using IBM SPSS Statistics (v.29). Prior to analyses, data were screened for missing values and fit of distributions with assumptions underlying univariate analysis. This included the Shapiro-Wilk test on z-score-transformed data and qqplot investigations for normality, Levene’s test for homogeneity of variance, and boxplot and z-score (±3.29) investigation for identification of influential outliers. Metrics of the central tendencies and variability were computed for each measure. Welch’s t-test was used to compare treatment groups during probe trials. Analysis of variance (ANOVA), including repeated measures, were used to analyze data where appropriate, and simple main effects were used to dissect significant interactions. Sex was included as a biological variable in all analyses. Where appropriate, the Greenhouse-Geisser or Huynh-Feldt adjustment was used to protect against violations of sphericity for repeated measures designs. Multiple pairwise comparisons were subjected to Bonferroni correction, where appropriate. Sex × genotype effects are reported where significant, otherwise data are discussed and visualized collapsed for sex. The critical alpha value for all analyses was p < 0.05 unless otherwise stated.

---

**Supplemental Figure 1:**
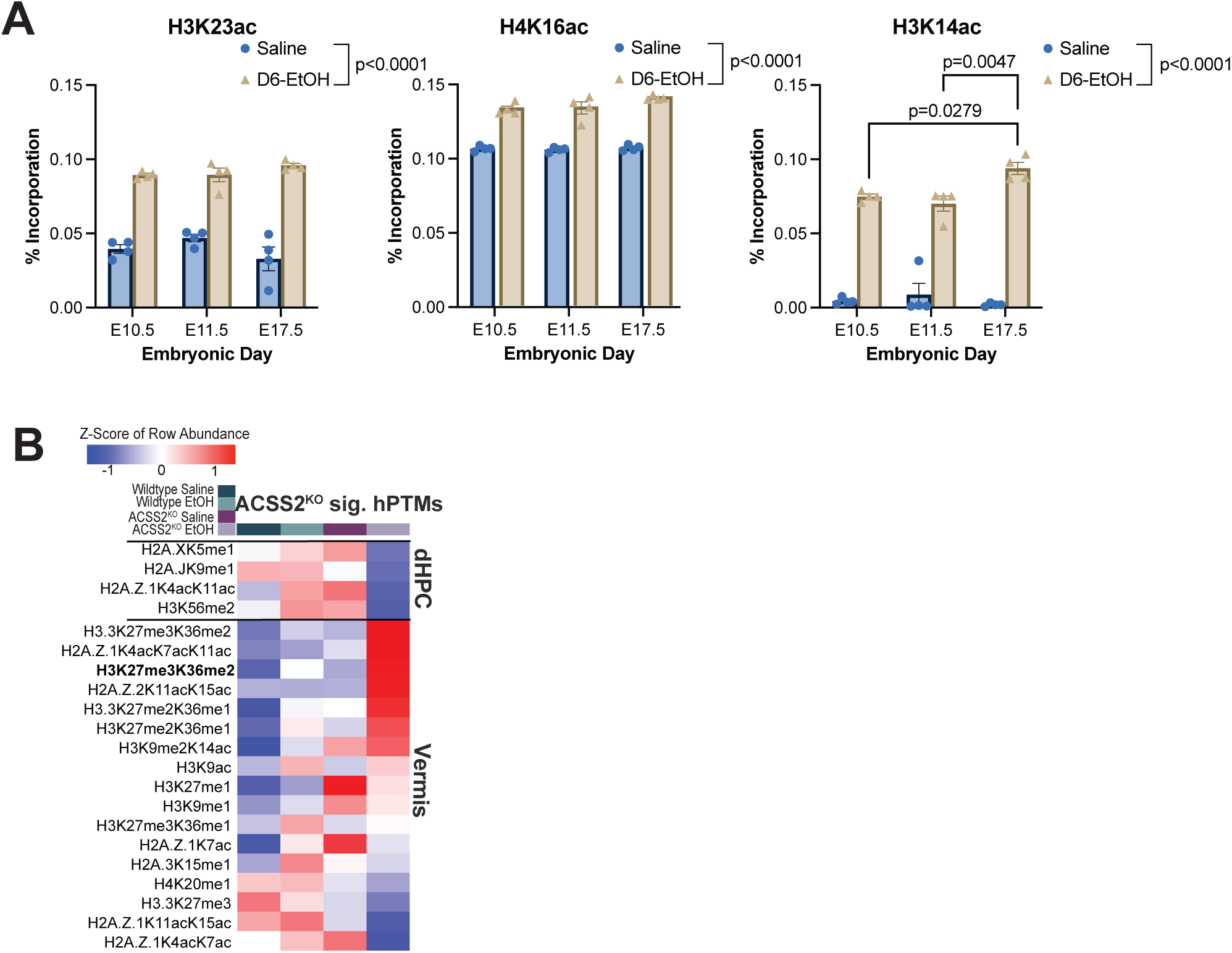
Prenatal alcohol exposure-related histone modifications. **(A)** Average percentage of heavy labeling incorporation into fetal brain histone peptides, H3K23ac (left), H4K16ac (middle), and H3K14ac (right) at embryonic timepoints E10.5, E11.5, and E17.5 following Saline or D6-EtOH exposure. Sample size n=4 per timepoint per condition. **(B)** Heatmap of average row Z-score of relative abundance for significant ACSS2^KO^ hPTMs sorted from most upregulated to most downregulated following cPAE in the dHPC (upper) and vermis (lower). Overlapping hPTMs are bolded. Sample size n=4 per timepoint per condition.

**Supplemental Figure 2.**
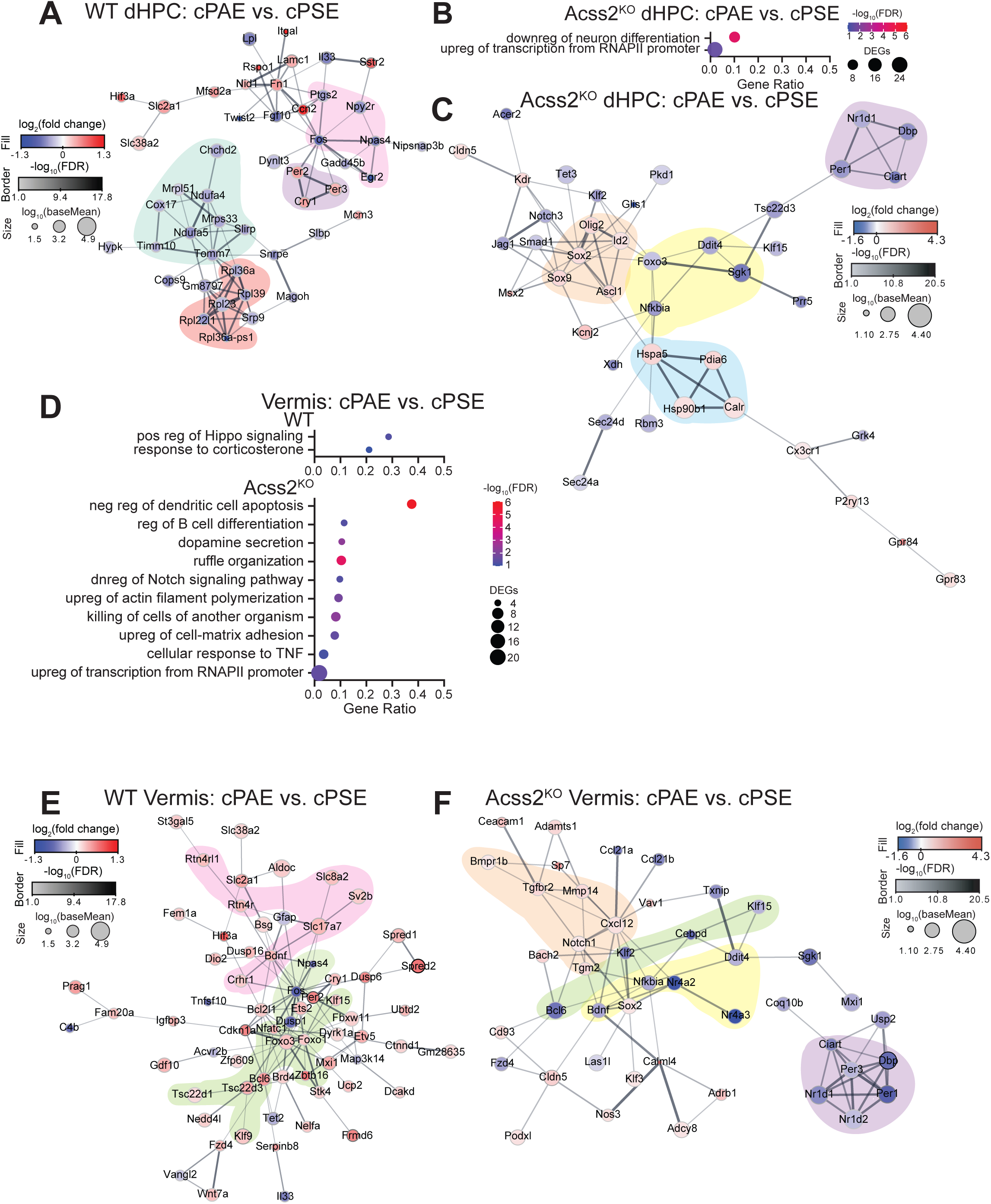
Functional analysis of cPAE-induced transcriptional changes in dHPC and vermis. **(A)** Protein-protein interaction (PPI) network diagram of DEGs in the WT cPAE dHPC.**B)** Significantly enriched GO terms in the ACSS2^KO^ cPAE dHPC. **(C)** PPI network diagram of DEGs in the ACSS2^KO^ cPAE dHPC. **(D)** Significantly enriched GO terms in the vermis of WT (upper) and ACSS2^KO^ (lower) cPAE mice. **(E)** PPI network diagram of DEGs in the WT cPAE vermis. **(F)** PPI network diagram of DEGs in the ACSS2^KO^ cPAE vermis. For PPI network diagrams, node fill color represents log_2_(fold change), node border color -log_10_(FDR), node size log_10_(baseMean), edge thickness STRING composite score. Sample sizes are n=4 per genotype per condition.

**Supplemental Figure 3.**
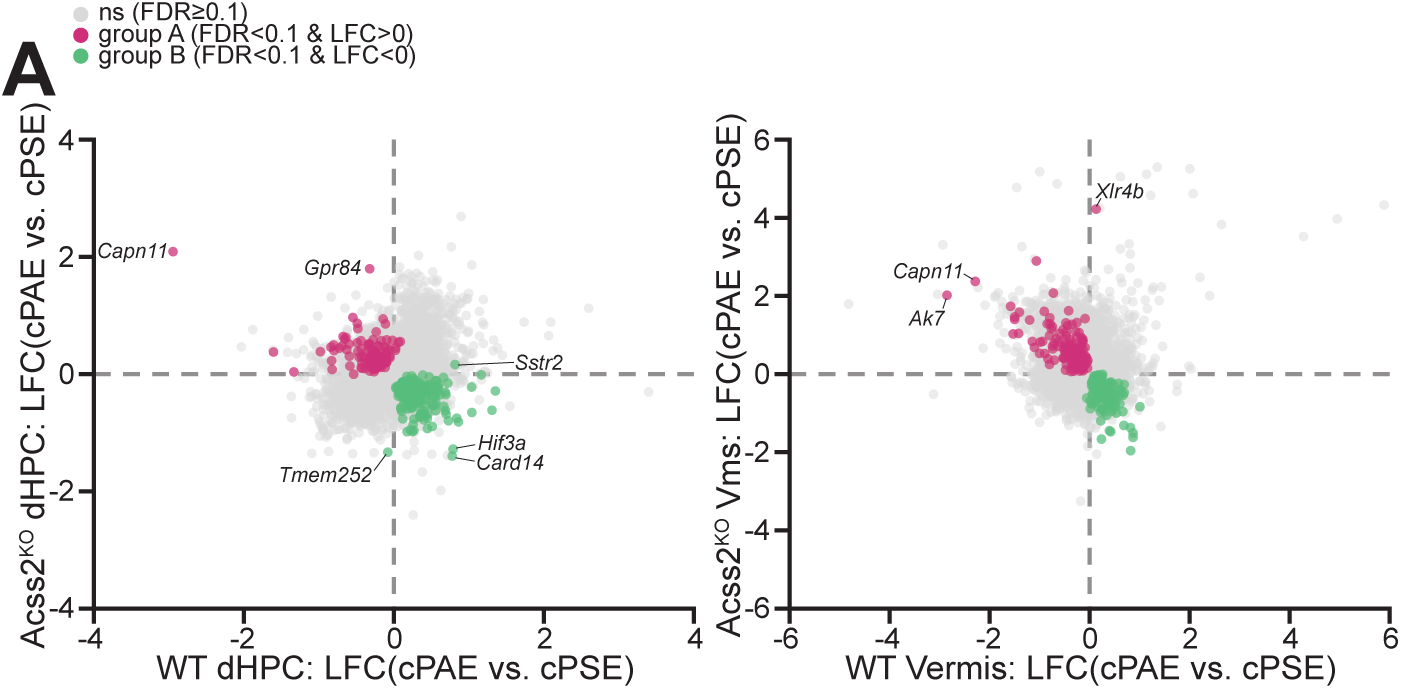
Genotype × exposure interaction genes in dHPC and vermis. **(A)** Scatterplots of gene level log_2_(fold change: cPAE vs. cPSE) comparing genotypes within each region. Dots represent genes (gray not significant, magenta Group A, green Group B). n=4 mice per genotype per condition.

**Supplemental Figure 4.**
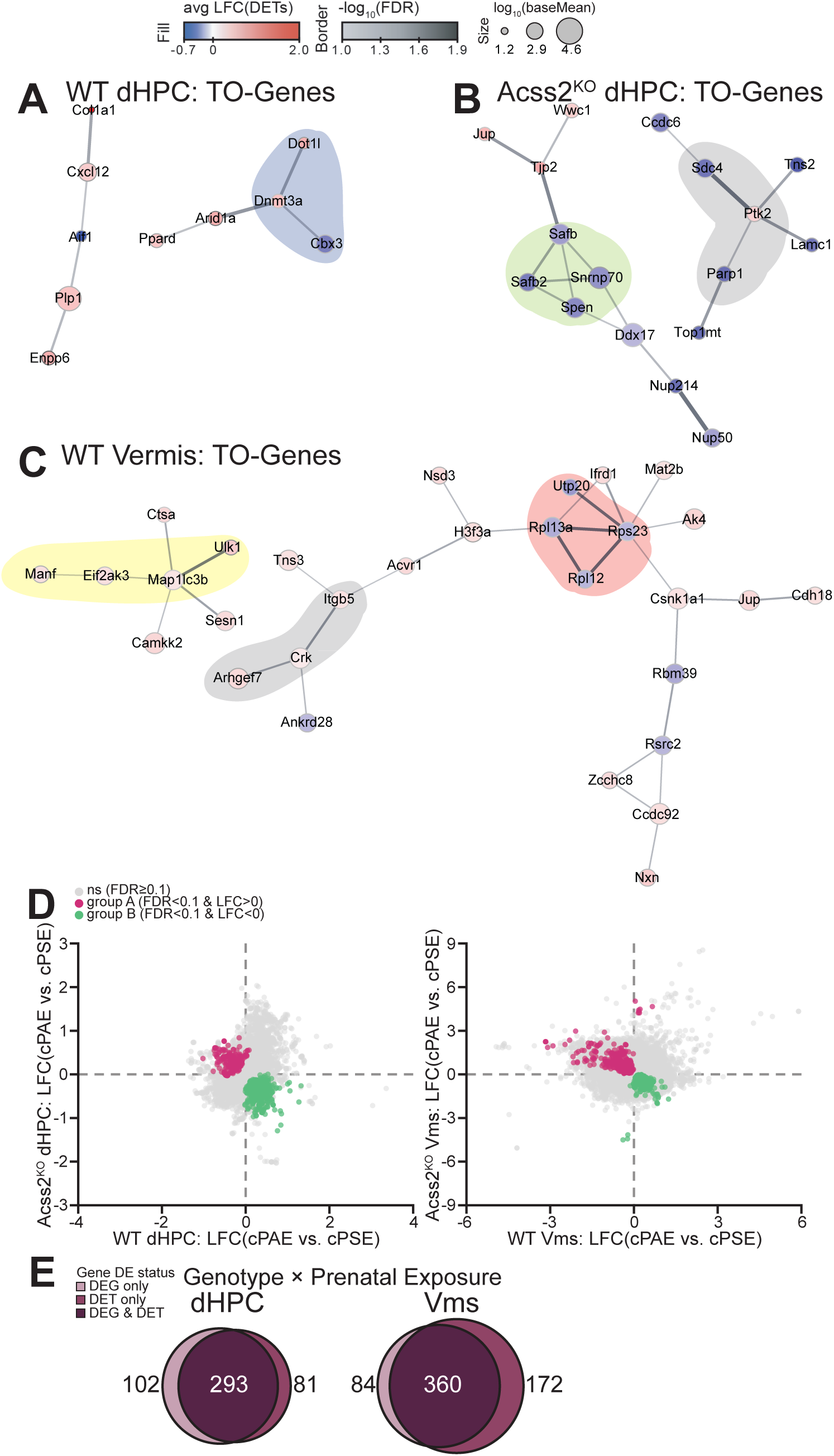
Functional analysis and genotype × exposure interaction analysis of genes differentially expressed only at the transcript level. **(A-C)** PPI network diagrams of TO-genes in the **(A)** WT cPAE dHPC, **(B)** ACSS2^KO^ cPAE dHPC, and **(C)** WT cPAE vermis. Node fill color and border color represent averages of log_2_(fold change) or -log_10_(FDR), respectively, across DETs encoded by the corresponding gene. Highlighting shows functional groups of interest. **(D)** Scatterplots of transcript level log_2_(fold change: cPAE vs. cPSE) comparing genotypes within each region. Dots represent transcripts (gray not significant, magenta Group A, green Group B). **(E)** Venn diagrams showing overlap of interaction genes between gene and transcript levels in the dHPC (left) and vermis (right). n=4 mice per genotype per condition.

**Supplemental Figure 5:**
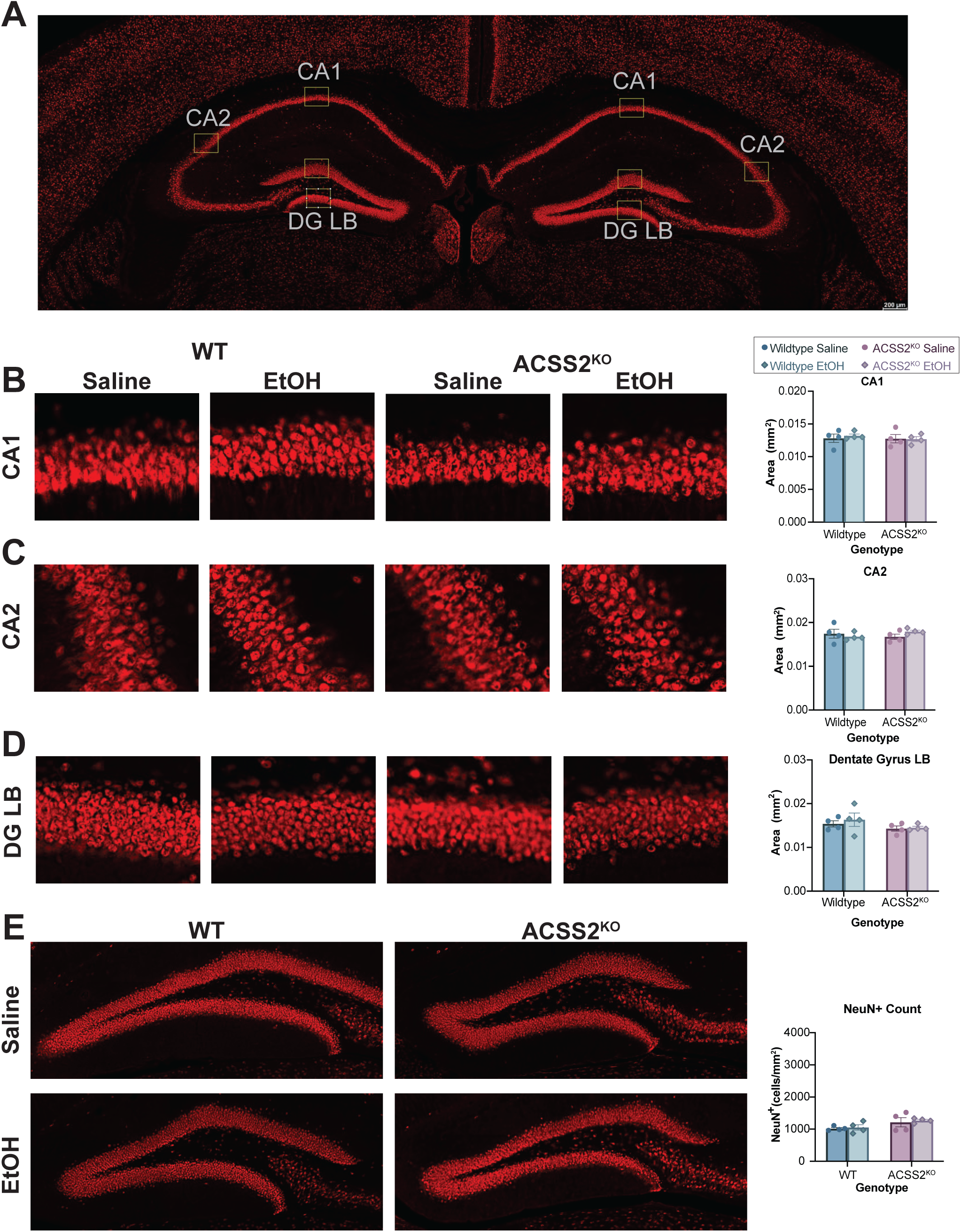
ACSS2 has no effect on female hippocampal thickness. **(A)** Hippocampal regions CA1, CA2, DG lower blade, and DG upper blade that were quantified for neuron thickness. **(B)** NeuN+ staining of hippocampal region CA1 from left to right in WT cPSE, WT cPAE, ACSS2^KO^ cPSE, and ACSS2^KO^ cPAE and the average thickness of CA1 region (right most) in females. **(C)** NeuN+ staining of hippocampal region CA2 from left to right in WT cPSE, WT cPAE, ACSS2^KO^ cPSE, and ACSS2^KO^ cPAE and the average thickness of CA2 region (right most) in females. **(D)** NeuN+ staining of dentate gyrus lower blade from left to right in WT cPSE, WT cPAE, ACSS2^KO^ cPSE, and ACSS2^KO^ cPAE and the average thickness of DG lower blade region (right most) in females. **(E)** Neun+ staining in the hilar region of WT cPSE (upper left), ACSS2KO cPSE (upper right), WT cPAE (lower left), and ACSS2KO cPAE (lower right) and average abundance of neurons (right most) in females. Sample size n=4 biological replicates per sex per condition per genotype.

**Supplemental Figure 6:**
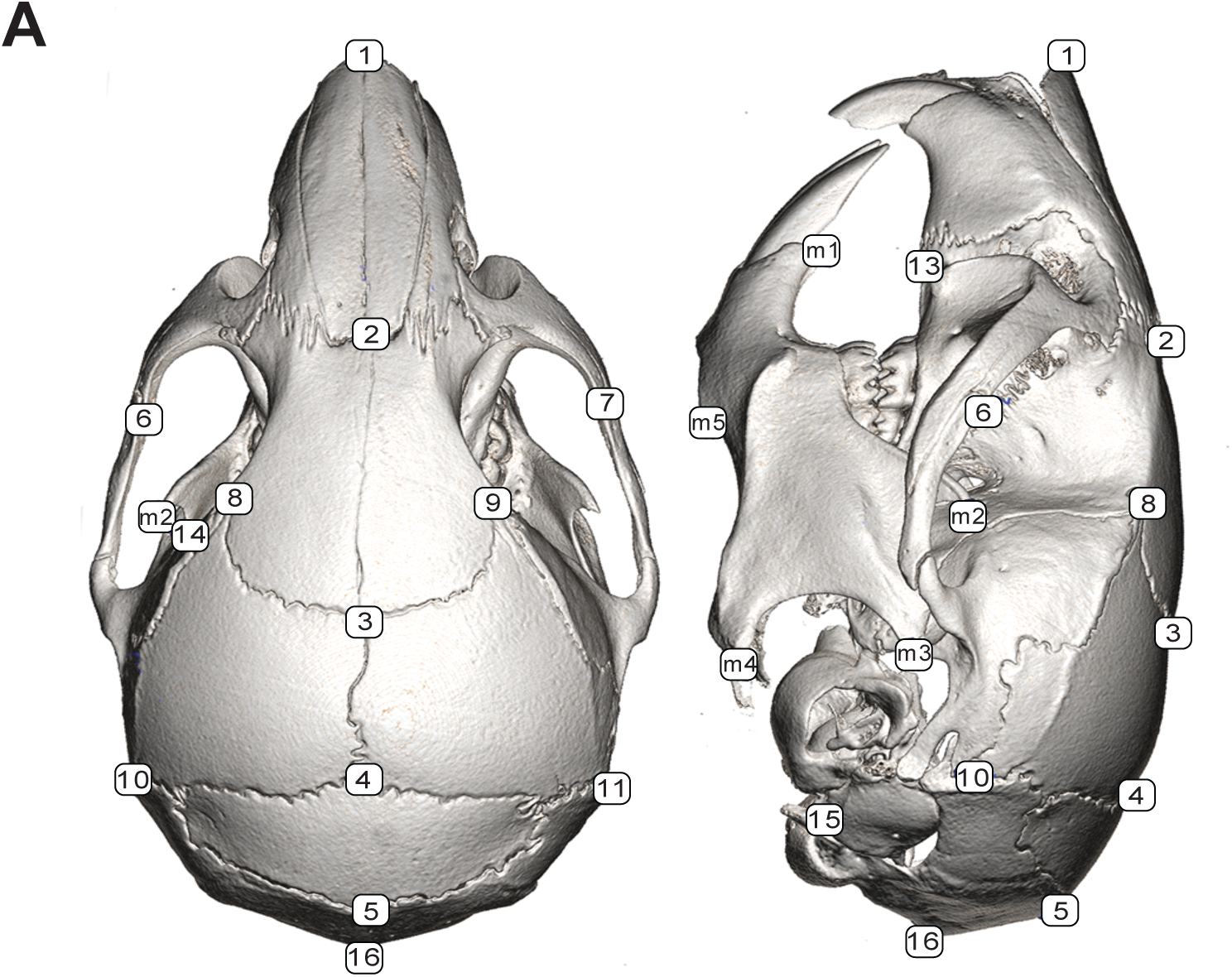
Craniofacial landmarks. **(A)** 21 skull and mandible landmarks collected from 3D reconstructions of CT scans.

**Supplemental Figure 7:**
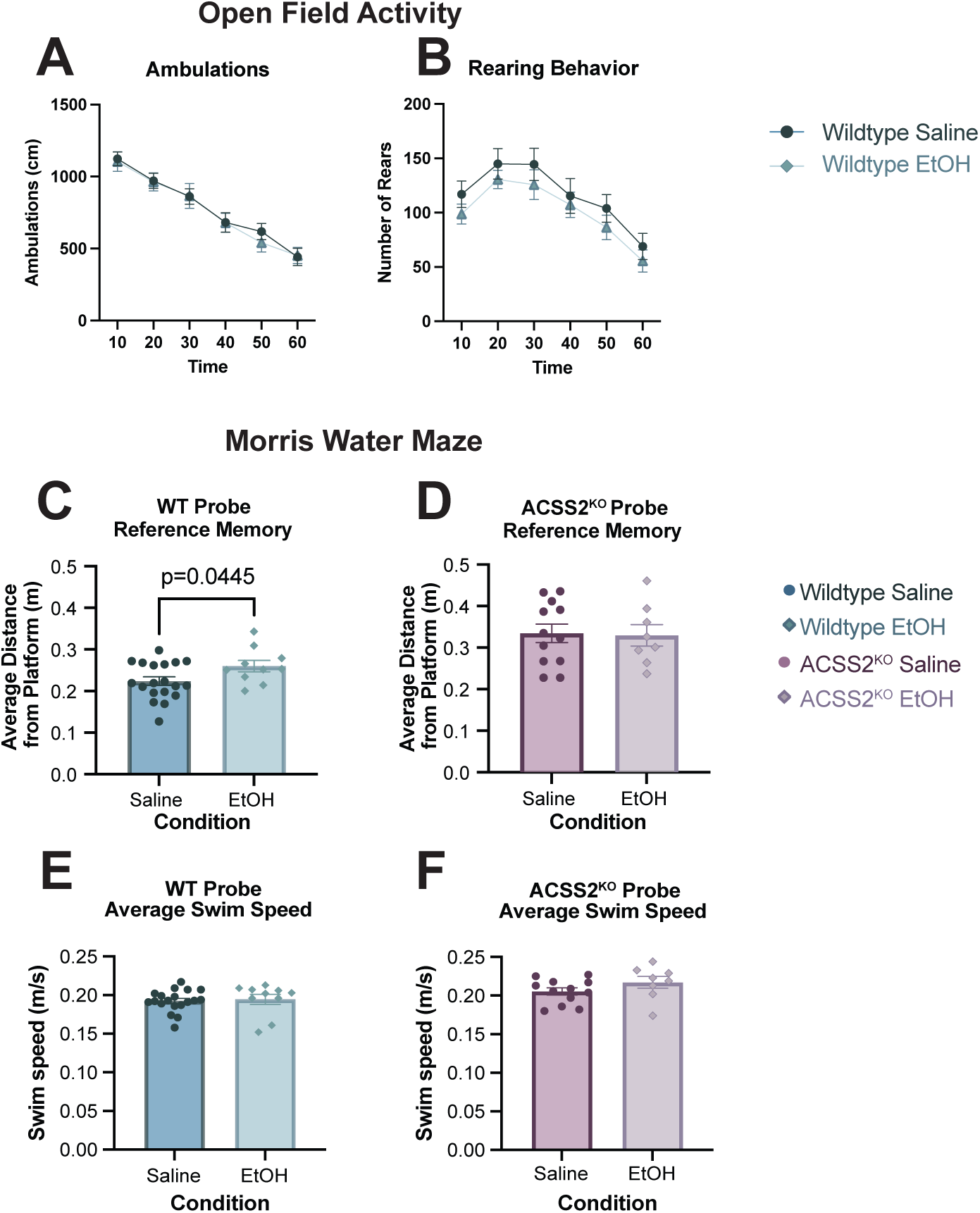
No changes in open field locomotor activity following cPAE in WT mice. **(A)** Average distance traveled (ambulation) across an hour during the open field activity. **(B)** Average number of rears across an hour during the open field activity. **(C)** Average distance WT mice were from platform during the probe trial. **(D)** Average distance ACSS2^KO^ mice were from platform during the probe trial. **(E)** Average swim speed of WT mice during the probe trial. **(F)** Average swim speed of ACSS2^KO^ mice during the probe trial. WT mice Sample size n=19 WT cPSE and n=10 WT cPAE, n=12 ACSS2^KO^ cPSE, and n=8 ACSS2^KO^ cPAE.

